# TSP1/TGF-β1 drives arachidonic acid metabolism to orchestrate neutrophil swarming

**DOI:** 10.1101/2025.11.25.688397

**Authors:** Lars Widera, Philippa Spangenberg, Devon Siemes, Sebastian Bessler, Jenny Bottek, Hannah Voss, Angelo Novak, Darleen Hueser, Mathis Richter, Alexander Potthoff, Bente Siebels, Manuela Moritz, Antonia Gocke, Stephanie Thiebes, Stephanie Tautges, Triantafyllos Chavakis, Jadwiga Jablonska, Tim Lämmermann, Matthias Gunzer, Hartmut Schlüter, Jan Hahn, Anika Grüneboom, Oliver Soehnlein, Klaus Dreisewerd, Jens Soltwisch, Olga Shevchuk, Daniel R. Engel

**Affiliations:** University Hospital Essen, Institute of Experimental Immunology and Imaging, Department Immunodynamics, Essen, Germany; University of Münster, Institute of Hygiene, Department of Biomedical Mass Spectrometry, Münster, Germany; Leibniz-Institut für Analytische Wissenschaften - ISAS-e.V., Department of Bioimaging, Dortmund, Germany; University of Münster, Institute of Experimental Pathology, Münster, Germany; University Medical Center Hamburg-Eppendorf, Section Mass Spectrometry and Proteomics, Center for Diagnostics, Hamburg, Germany; University Medical Center Hamburg Eppendorf, Center for Molecular Neurobiology Hamburg (ZMNH), Hamburg, Germany; University Hospital Dresden, Institute for Clinical Chemistry and Laboratory Medicine, Dresden, Germany; University Hospital Essen, Department of Otorhinolaryngology, Head and Neck Surgery, Essen, Germany; University of Münster, Institute of Medical Biochemistry, Center for Molecular Biology of Inflammation (ZMBE), Münster, Germany; University Hospital Essen, Institute of Experimental Immunology and Imaging, Essen, Germany; University Medical Center Hamburg-Eppendorf, Mildred Scheel Cancer Carrer Center Hamburg, Hamburg, Germany

## Abstract

Neutrophil swarming has emerged as a conserved multicellular behaviour observed across tissues and pathological contexts. Yet, the molecular cues and the spatially coordinated cellular circuits that drive the process of neutrophil swarming leading to cluster formation remain poorly understood. Here, we combine spatial proteomics and lipid profiling in a model of urinary tract infection to define the epithelial-immune circuits driving neutrophil cluster formation. We identify thrombospondin-1 (TSP1)-mediated activation of transforming growth factor beta 1 (TGF-β1) as key epithelial signal licensing neutrophil clustering and enhancing bacterial control. Spatial lipid analysis further reveals that TSP1/TGF-β1 signalling locally activates arachidonic acid metabolism in epithelial neutrophils, with 5-lipoxygenase dependent leukotriene synthesis required for swarm formation and infection clearance. These findings uncover a spatially coordinated defence mechanism in which epithelial-derived TSP1/TGF-β1 engages neutrophil lipid metabolism to orchestrate neutrophil swarming behaviour and reinforce antibacterial immunity.

Graphical abstract

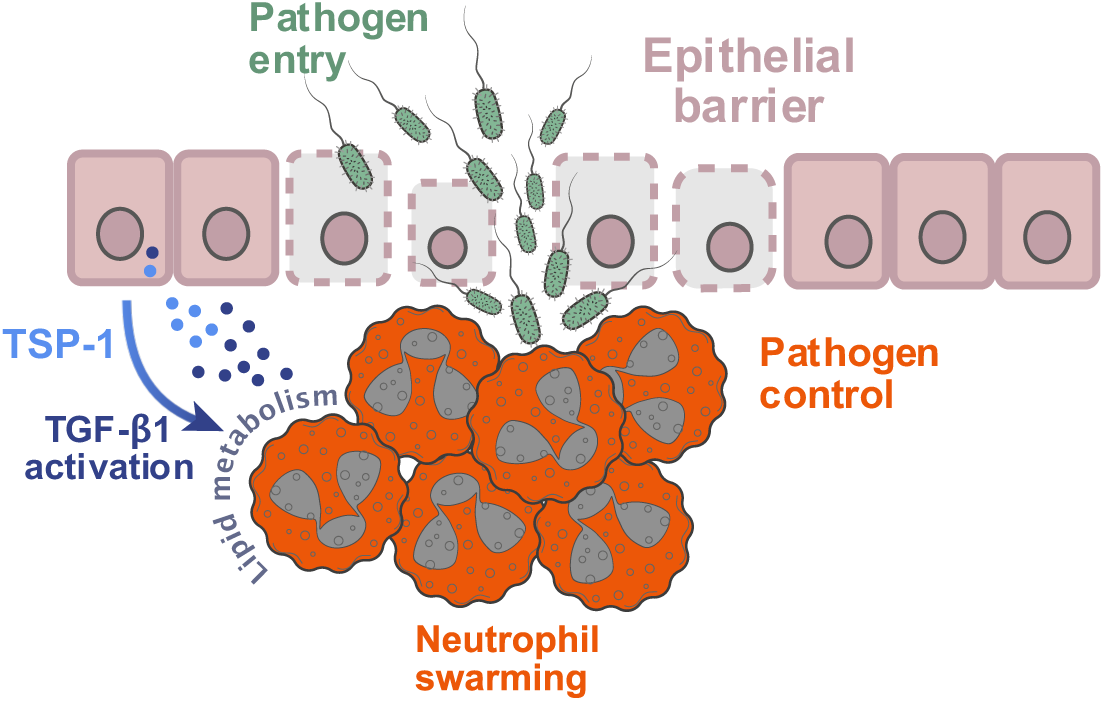

## Introduction

Epithelial surfaces are the bodýs frontline defence, serving as both a physical barrier and an active regulator against microbial entry.^1,2^ Once breached, pathogens trigger the rapid recruitment of neutrophils, which self-organize into multicellular clusters that contain infection and prevent systemic spread.^3–5^ A hallmark of this process is ‘neutrophil swarming’, a collective migratory behaviour mediating dense neutrophil clusters at sites of microbial challenge.^6–8^ Such local accumulations are observed in diverse contexts, yet the molecular circuits that coordinate neutrophil swarming during infection remain poorly defined.

Neutrophil swarm formation unfolds in distinct stages, starting with pioneer neutrophils responding to danger-associated molecular patterns and chemotactic mediators,^9,10^ followed by an amplification loop through secondary cell death.^7,11^ Central to this amplification is the arachidonic acid (AA) metabolism leading to leukotriene B_4_ (LTB_4_) synthesis, which stabilizes chemotactic gradients and sustains directional migration.^7,12^ Termination occurs through chemokine receptor desensitization and conversion of LTB_4_ into antagonistic metabolites.^13,14^ Disruption of LTB_4_ receptor or the enzyme 5-lipoxygenase (Alox5), which is required for LTB_4_ synthesis, abolishes neutrophil swarming and cluster formation.^7,15^ LTB_4_ biosynthesis begins when cytosolic phospholipase A_2_ (cPLA_2_) liberates AA from membrane phospholipids, followed by conversion through Alox5 into leukotriene intermediates.^16^ While these mechanistic insights are well defined in sterile inflammation models,^7,17,18^ the molecular cues that regulate arachidonic acid metabolism and LTB_4_-driven swarming under *in vivo* infectious conditions remains poorly understood.

To address this gap, we employed a murine model of urinary tract infection (UTI) induced by uropathogenic *Escherichia coli* (UPEC). Acute UTI is considered the most prevalent bacterial infection worldwide, affecting more than 150 million individuals annually.^19–21^ Neutrophils constitute the first group of immune cells at the infected epithelium, and are typically detectable within two hours post infection.^22,23^ Their initial recruitment is initiated by chemokines secreted by resident macrophages and epithelial cells.^23–26^ Here, we identify epithelial thrombospondin-1 (TSP1)–mediated activation of TGF-β1 as a critical signal, which license neutrophil clustering and promote infection control. We show that this TSP1/TGF-β1 signalling licences AA metabolism in epithelial neutrophils, and that pharmacological inhibition of the AA metabolism disrupts neutrophil clustering and infection control. These findings define the epithelial-neutrophil communication pathway that reinforces epithelial defence through localized lipid-mediator signalling.

## Results

Epithelial surfaces are constantly breached by pathogens. In this situation, the immunity of the epithelium is insufficient, causing neutrophils to migrate into the infected epithelium, form local clusters and fight the infection. Using a murine model of bacterial urinary tract infection, we visualised the spatial accumulation of neutrophils *in situ* with 3D light-sheet fluorescence microscopy (Figure 1A, B). To study the microenvironmental signals driving neutrophil cluster formation, we performed LC-MS/MS-based proteomic profiling of UPEC-infected and control bladders (Fig. 1C). Principal Component Analysis (PCA) demonstrated a distinct segregation of the proteome in infected over control samples along the 1^st^ principal component, accounting for 27% of the explained variance (Fig. 1D). In total, 6,904 proteins were identified, with 173 significantly upregulated and 75 downregulated proteins registered in UPEC-infected bladders compared to healthy controls (Fig. 1E, Table S1). STRING network analysis in combination with an unsupervised DBSCAN clustering of the 173 upregulated proteins identified 12 distinct clusters, with the largest cluster comprising of 18 proteins for the pathway “Leukocyte cell-cell adhesion” (Fig. 1F, Table S2 and Table S3). Within this cluster, Thrombospondin-1 (TSP1) emerged as an important candidate due to its dual role in mediating cellular adhesion and modulating inflammatory signalling, and strong interaction with the proteins of the largest cluster (Fig. 1G). Next, we confirmed the secretion of TSP1 protein within the infected bladder tissue using ELISA (Fig. 1H), immunofluorescence microscopy (Fig. 1I) and mRNA *in situ* hybridization (Fig. 1J). Notably, *Tsp1* and *Tgf-β1* – the latter being an important interaction partner of TSP1 – colocalized within epithelial regions, indicating coordinated local expression (Fig. 1J).

**Figure 1.**
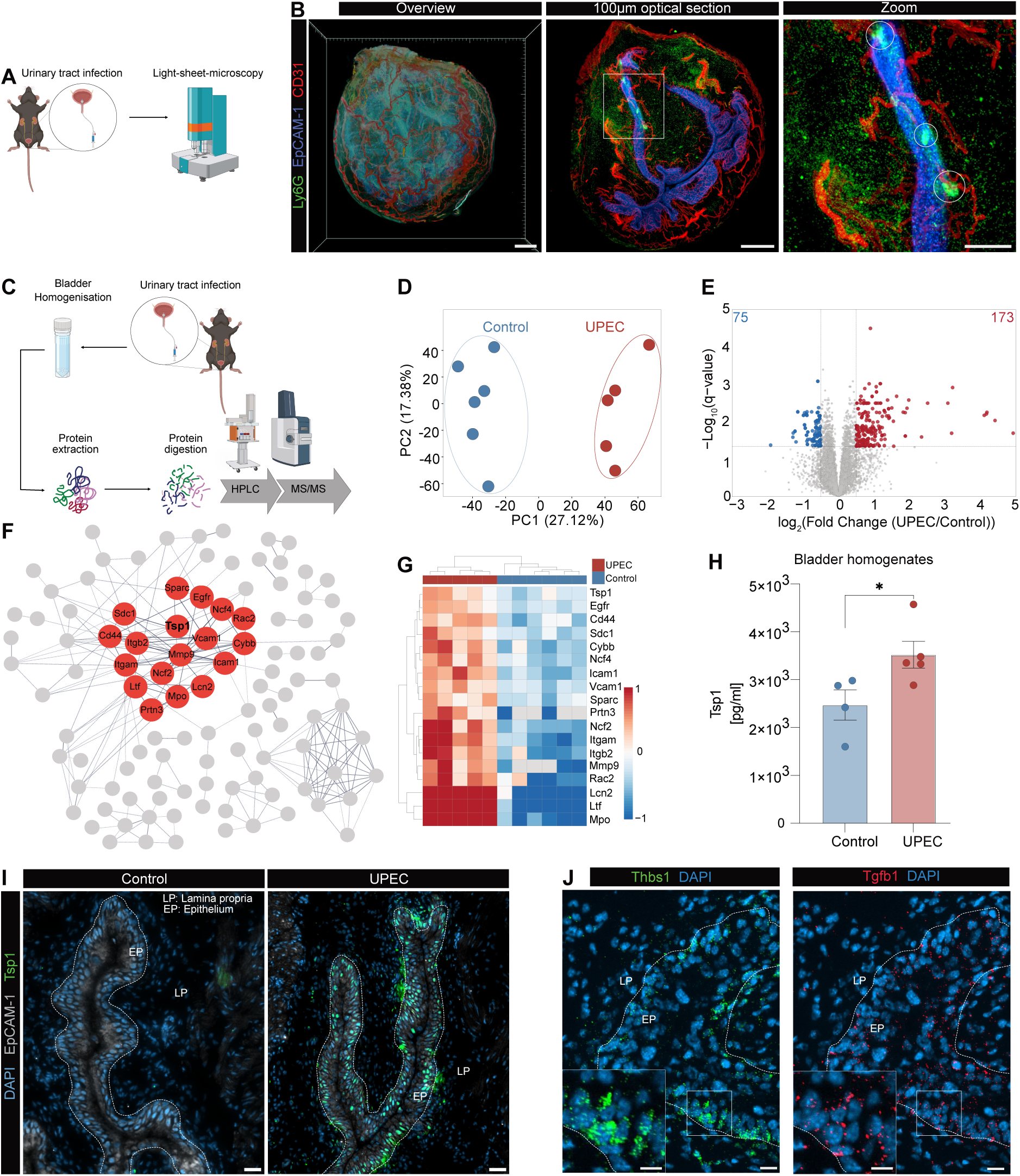
Neutrophil cluster formation is associated with the TSP1/TGF-β1 axis as response to UPEC infection. **(A)** Experimental overview of 3D light sheet microscopy performed 24 hours after UPEC infection. Partially created in BioRender. Widera, L. (2025) https://BioRender.com/bb0zujs. **(B)** Light sheet microscopy of the whole urinary bladder 24 hours after infection. Antibodies against Epcam-1 (blue), Ly6G (green), CD31 (red) were injected 4 hours prior sacrifice. The scale bar indicates 500 µm (overview) and 200 µm (zoom). **(C)** Experimental overview of the proteomic LC-MS/MS workflow performed 24 hours after UPEC infection. Partially created in BioRender. Widera, L. (2025) https://BioRender.com/bb0zujs **(D)** Dimensionality reduction of the LC-MS/MS data of all identified proteins (n=6904). Scatter plot represents the principal components (PC) 1 and 2 from nonlinear iterative squares PC Analysis (PCA). **I** The significantly regulated lipids (*q* value < 0.05 and |log_2_(FC)| > 0.5) are coloured in the Volcano plot (173 upregulated and 75 downregulated). **(F)** STRING Protein-Protein interaction network of upregulated proteins, in UPEC condition (q-value < 0.05; Log_2_ (FC > 0.5). Clustering by DBSCAN. The proteins of the largest cluster (cluster in red, n=18 proteins; Table S2) were found to be associated with “Leukocyte cell-cell adhesion” (Table S3). Line thickness indicates the confidence of interactions with a minimum required interaction score >0.7. Disconnected nodes were hidden to improve visualisation clarity. **(G)** Relative protein abundance of proteins of the “leukocyte cell–cell adhesion” cluster (shown in F) upon UPEC infection. **(H)** Expression of TSP1 protein levels in bladder homogenates by ELISA 24 hours after UPEC infection. **(I)** Immunofluorescence microscopy of bladder tissue from control (n=3) and UPEC-infected mice (n=6) 24 hours after infection. The scale bar indicates 20µm. **(J)** Fluorescence *in situ* hybridisation for the detection of Thbs1 (green) and TGF-β1 mRNA (red) in UPEC-infected bladder tissue. The scale bar indicates 20 µm (overview) and 10 µm (zoom), respectively. (n=3). Dotted lines indicate the epithelium stained by Epcam-1. *p<0.05, Mann-Whitney; Data are mean±SEM; EP=epithelium, LP=lamina propria).

Activation of TGF-β1 is mediated by TSP1-dependent release of TGF-β1 from its latency-associated peptide (LAP). To study the role of TSP1-mediated TGF-β1 activation, we used the mimetic peptide LSKL that selectively blocks the TSP1-domain responsible for TGF-β1 activation. To examine the role of TSP1/TGF-β1, we determined neutrophil localization and abundance in bladder tissue sections 6 and 24 hours post infection. In the absence of infection, neither PBS nor TGF-β1 inhibitor-treated mice displayed neutrophil infiltration (Fig. 2A). Following UPEC infection, neutrophils progressively accumulated within the epithelium after 6 and 24 hours post-infection, accompanied by decline in the underlying lamina propria (Fig. 2A, B), suggesting spatial redistribution of neutrophils from the lamina propria to the epithelium. By contrast, inhibition of the TSP1/TGF-β1 axis significantly prevented epithelial redistribution, while neutrophil numbers in the lamina propria remained unchanged (Fig. 2A, B). This defect correlated with an increased bacterial burden in the bladder at 24 hours (Fig. 2C), indicating that TSP1-mediated TGF-β1 activation is required for effective neutrophil accumulation into the infected epithelium and infection control. To systematically quantify clustering of neutrophils in the infected epithelium, we developed a custom image analysis pipeline, that computes tissue-morphology-aware pairwise cell-cell distances and identifies clusters using the Markov Cluster algorithm. (Fig. 2D). Large neutrophil clusters (>100 neutrophils) formed exclusively in UPEC-infected bladders and were abrogated by TSP1/TGF-β1 inhibition (Fig. 2D, E). Together, these data demonstrate that TSP1/TGF-β1 signalling promotes high-density neutrophil cluster formation in the infected epithelium.

**Figure 2.**
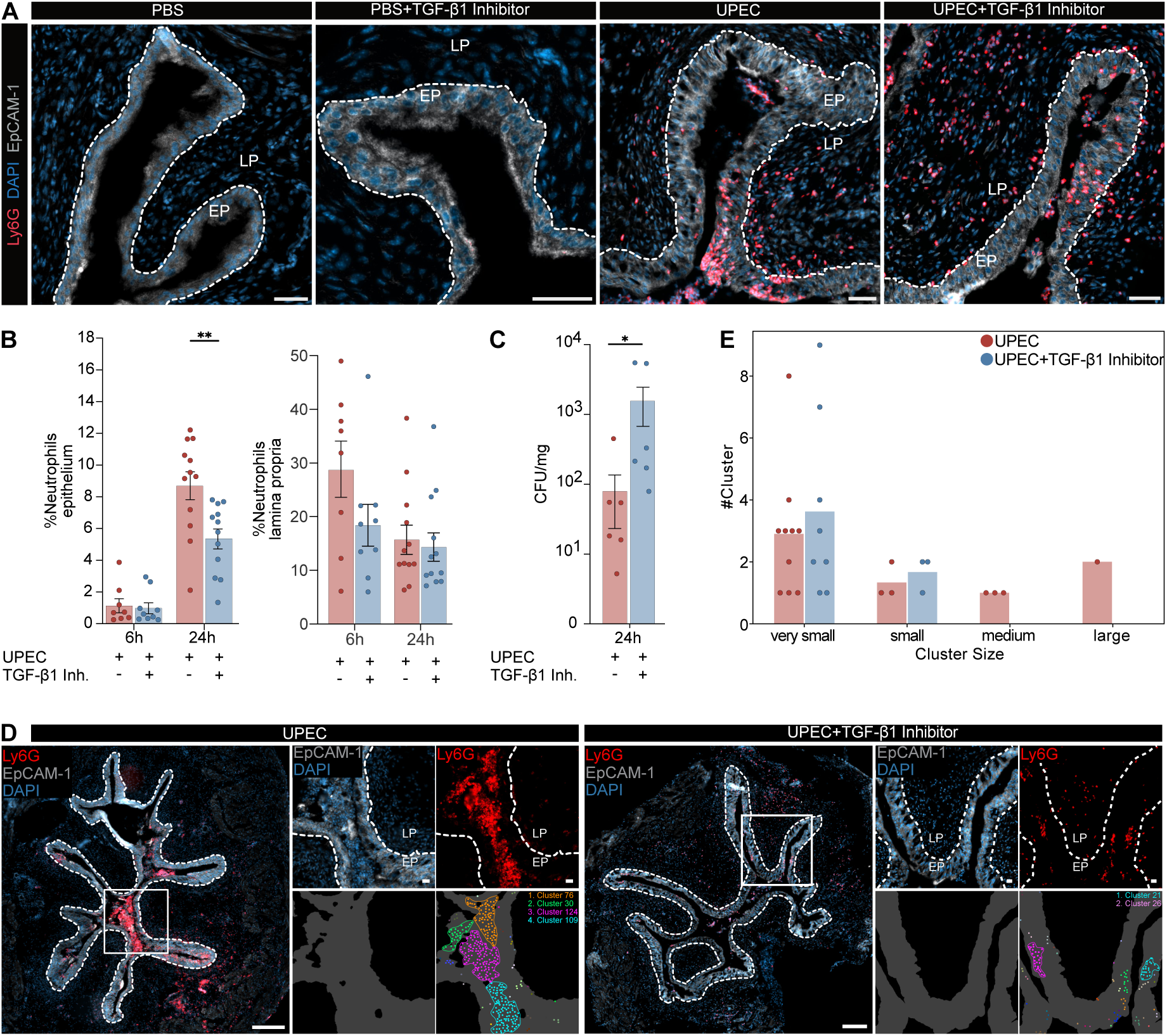
TSP1-dependent TGF-β1-release stimulates neutrophil infiltration and clusters in the epithelium. **(A)** Bladder tissue sections were stained with Ly6G (neutrophils), EpCAM-1 (epithelial cells) and DAPI (nucleus) 24 hours after infection. Scale bar indicates 100 µm. **(B)** Quantitative analysis of neutrophil infiltration shown in of (A). **(C)** Quantification of bacterial burden using colony forming units (CFU) of bladder homogenates 24 hours post-infection. **(D)** Immunofluorescence images of bladder tissue 24 hours after UPEC infection, stained for EpCAM-1 (grey), Ly6G (red), and DAPI (blue). Lower panels show algorithm-based cluster segmentation with color-coded clusters. Scalebar indicates 200 µm (overview) and 50 µm (zoom). **(E)** Quantitative results of the images shown in D (very small=10-30, small=30-50, medium=50-100, large >100 neutrophils/cluster). *p<0.05, **p<0.01 Mann-Whitney; Data are mean±SEM; EP=epithelium, LP=lamina propria.

To evaluate neutrophil clustering within the intact 3D tissue architecture, we employed light-sheet fluorescence microscopy, enabling high-resolution, volumetric imaging of the entire organ. This approach allowed comprehensive, whole organ visualization and quantification of neutrophil clusters upon TSP1/TGF-β1 inhibition (Fig. 3A). Quantitative analysis revealed a reduction in both number and size of neutrophil clusters within the infected epithelium following TSP1/TGF-β1 inhibition (Fig. 3B to D). A negative binomial regression model revealed higher expected number of clusters in UPEC bladders compared to TSP1/TGF-β1 inhibitor–treated bladders, with stable/slowly-decreasing differences across cluster size distributions (Fig. 3E, F and Table S4).

**Figure 3.**
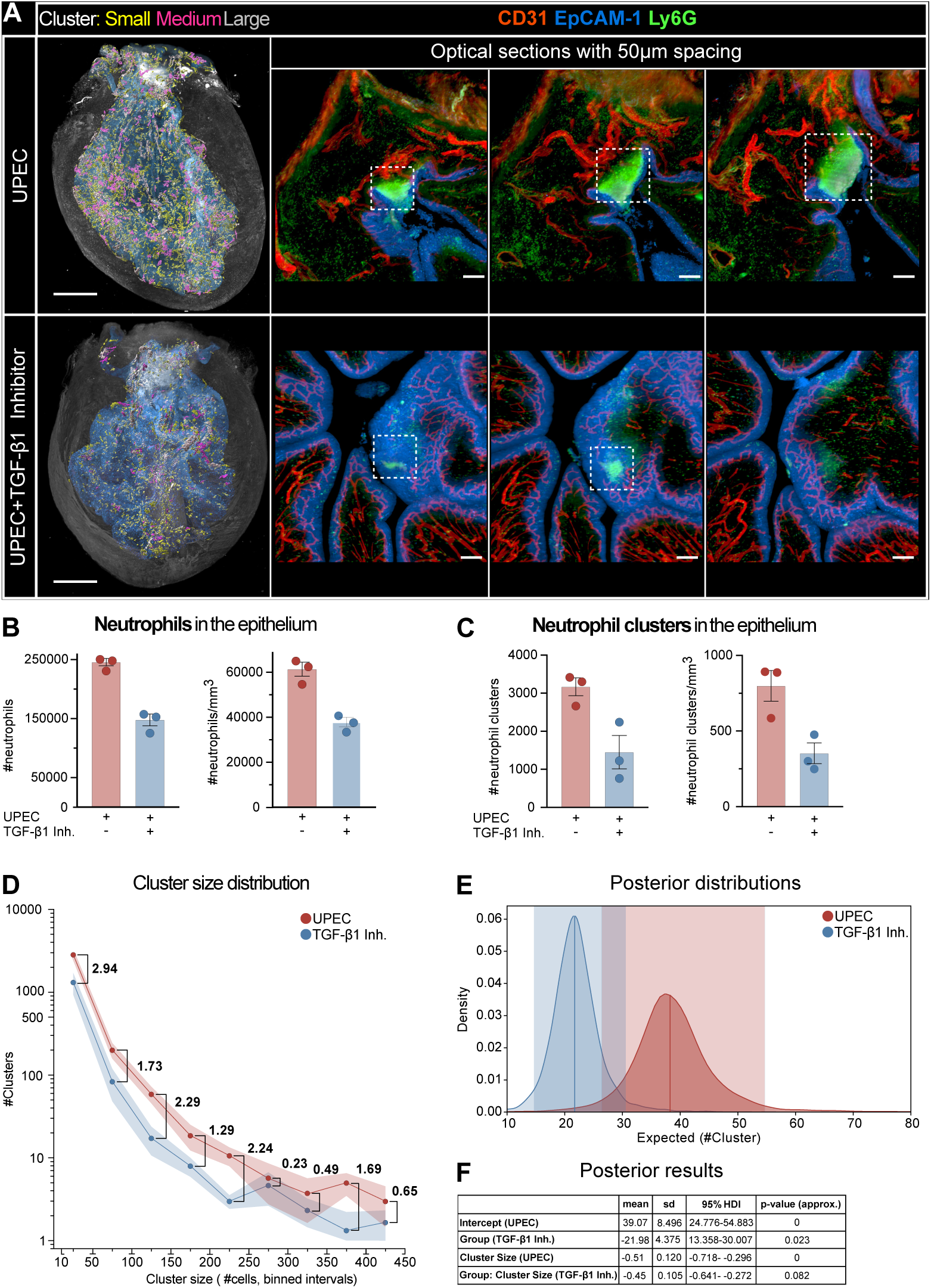
3D imaging reveals reduced neutrophil clusters in the absence of TGF-β1. **(A)** Light sheet fluorescence microscopy images of whole bladders 24 hours after infection, stained with Ly6G (neutrophils), EpCAM-1 (epithelial cells) and CD31 (blood vessels). The overview images to the left show the epithelium (blue) and neutrophil clusters coloured regarding cluster size (small=10-50 (yellow), medium=50-100 (pink), large>100 (grey) neutrophils. Scalebar indicates 300µm. The three detailed images show Z-stacks with 50µm thick optical sectioning. Scalebar indicates 100µm. **(B)** Quantitative results of neutrophil infiltration in the epithelium shown in (A). **(C)** Quantitative results of neutrophil cluster formation in the epithelium shown in (A). **(D)** Quantitative results of neutrophil clusters, showing the count of identified clusters for each cluster size bin. Numbers indicate the Coheńs effect size. **(E)** Exponentiated posterior distributions of expected cluster counts from a Bayesian negative binomial generalized linear mixed model (NB-GLMM). Shaded areas indicate 95% highest density intervals (HDI). **(F)** Posterior summaries of model parameters on the natural scale, showing mean, standard deviation (SD), and 95% HDI. Approximate one-sided p-values were derived from the log-scale posteriors.

We next asked whether neutrophil maturation could explain impaired clustering. Using a multiplex spectral flow cytometry panel including CD101, CD62L, CXCR2, CD16, Ly6G, and c-Kit, bladder neutrophils were defined as Ly6G⁺Ly6C⁺, with doublets and dead cells excluded (Fig. S4A). We observed that maturation profiles were largely unchanged between time points and treatment groups (Fig. S4B). Thus, TSP1/TGF-β1–dependent clustering is independent of neutrophil maturation. To study the underlying mechanism of neutrophil cluster formation, we performed 3D spatial tissue proteomics with the Nano-second-Infrared Laser (NIRL) ablation technology (Fig. 4A). This technology enables layer-by-layer analysis of the epithelium, thereby revealing the local mechanisms, and local production of TSP1, for neutrophil cluster formation. Ablated tissue regions were subjected to LC-MS/MS-based proteomics, enabling comparative 3D proteome profiling across the distinct tissue layers. In total, 4158 proteins were quantified across all laser-ablation layers (Table S5 to Table S8). Clustering of the different laser ablation layers revealed 6-8 spatially ordered clusters based on proteome profile similarity (Fig. 4B, Fig. S1 and Fig. S2). Comparison to published cell-layer-specific marker proteins of the bladder tissue architecture,^27^ enabled a spatially resolved proteomic map encompassing superficial, intermediate, and basal epithelial cells, the lamina propria, and the detrusor muscle (Fig. 4B). Among the identified molecules, TSP1 displayed distinct compartmental localization in both healthy and UPEC-infected bladders, with the highest abundance in basal epithelial and lamina propria cells (Fig. 4C). Correlation of the proteome profiles with TSP1 abundance (Pearson correlation coefficient > 0.7; p-value < 0.01) revealed abundance of neutrophil-associated proteins such as matrix metalloproteinase-9 (Mmp9), neutrophil elastase (Elane), and cathepsin G (Ctsg) under infectious, but not during steady state conditions (Fig. 4D, Fig. S3, Table S9). Functional enrichment revealed the involvement of these proteins in neutrophil migration and lipid metabolism under infectious conditions, whereas developmental and ECM-related processes were dominant in healthy tissue (Fig. 4E, Table S10 and Table S11). Moreover, TSP1-correlated proteins not only included neutrophil-specific molecules, but also enzymes of the arachidonic acid (AA) pathway, among them Alox5 and Alox5 activating protein (Alox5ap) (Fig. 4F). These data suggest that TSP1 promotes neutrophil-driven and lipid-mediated inflammation during infection.

**Figure 4.**
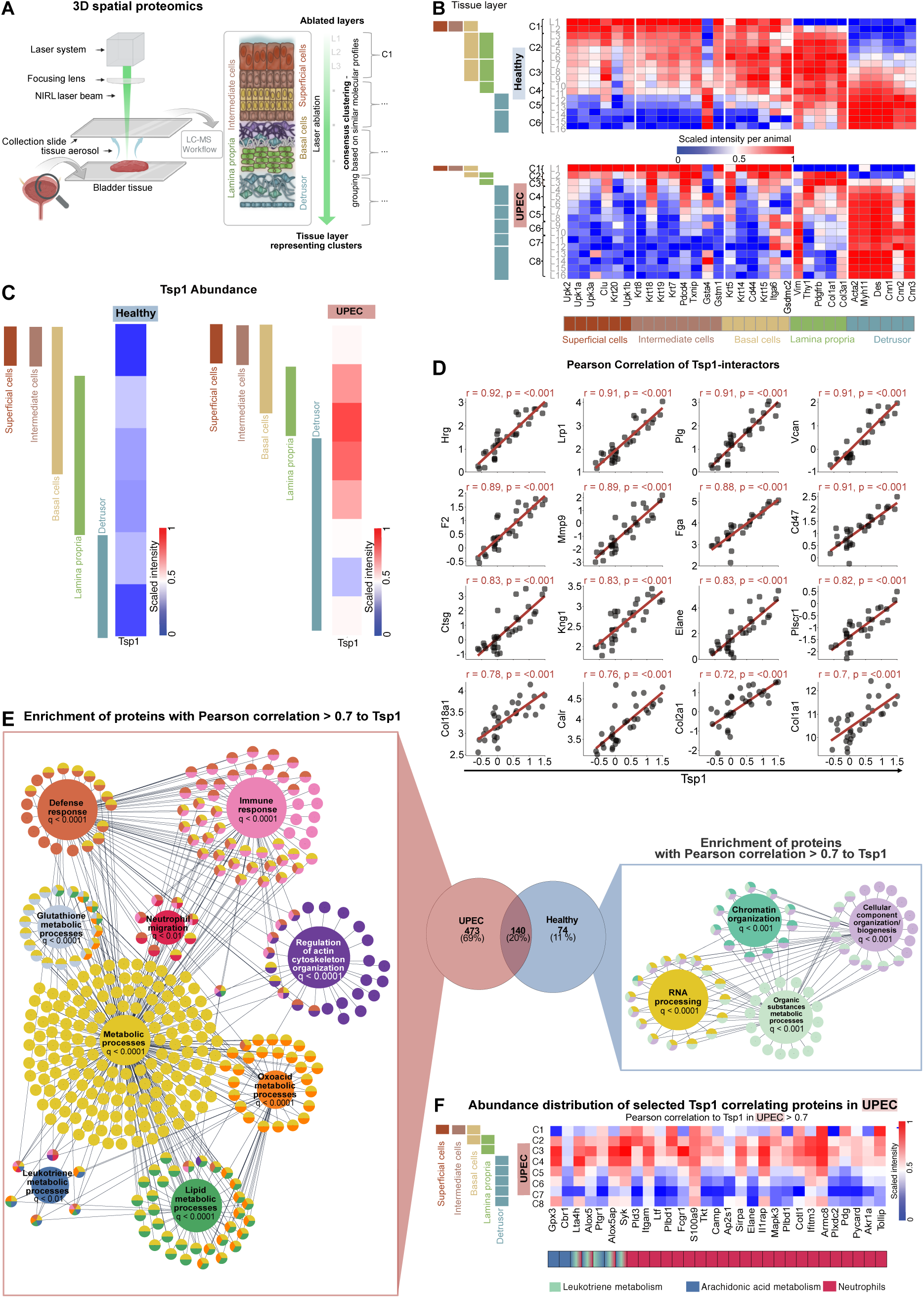
Spatial proteomics links TSP1 localization to neutrophil lipid metabolism. **(A)** Experimental overview of the NIRL-based tissue layer-resolved 3D proteome analysis of healthy (n=2) and UPEC infected (n=2) mouse bladders. Partially created in BioRender. Widera, L. (2025) https://BioRender.com/t86f70g. **(B)** Scaled abundance of tissue layer-specific markers^27^ across the laser ablation layers and proteome profiles. **(C)** Scaled abundance distribution of TSP1 across the different tissue layers. **(D)** Pearson correlation of known Tsp1 interactors.^53^ Proteins with linear correlation > 0.7 are highlighted. The Pearson correlation coefficient was calculated on log2 transformed, normalized, and imputed values. Scaling was not applied prior to correlation. **(E)** Venn diagram of proteins that correlate (r > 0.7) with Tsp1 in UPEC-infected and healthy bladders. Proteins that specifically correlate in healthy and UPEC-infected bladders, a STRING-based protein-protein interaction network was computed. Interactions were accepted at a high confidence > 0.8 and all interaction sources were used. Selected, significant (FDR < 0.05) GO-BP gene sets are shown and connected to their affiliated proteins. Singleton and unconnected proteins are hidden. (UPEC: 370/487 elements are shown; healthy: 33/63 elements are shown). Complete enrichment tables and Venn overlays can be found in (Table S10 and S11). **(F)** Abundance distribution of proteins involved in leukotriene metabolism, arachidonic acid metabolism and neutrophils with a Pearson correlation (r > 0.7) to Tsp1 abundance in UPEC infected but not healthy samples, across different tissue layers.

To further examine the role of lipid metabolism in neutrophil cluster formation, we employed matrix-assisted laser desorption/ionization mass spectrometry imaging (MALDI-MSI) in combination with immunofluorescence microscopy (Fig. 5A). Consecutive tissue sections were used to map the spatial distribution of molecular species by MALDI-MSI and to localize neutrophils by immunofluorescence staining, enabling spatial alignment of lipid features with neutrophil-rich regions (Fig. 5A). Co-registration of MALDI-MSI and microscopy data, combined with a machine-learning classifier, identified four molecular species that showed strong spatial colocalization with Ly6G⁺ neutrophils (Fig. 5A, B). These molecular species served as a proxy for neutrophil distribution in the MALDI-MSI dataset. Analysis of the neutrophil-specific MALDI-MSI profiles revealed two distinct lipid-defined neutrophil clusters (Fig. 5C). Mapping these clusters back onto the tissue demonstrated that they corresponded to neutrophils localized either within the lamina propria or within the epithelium (Fig. 5C). Following TSP1/TGF-β1 inhibition, 51 lipids were differentially regulated in epithelial neutrophils, whereas only 6 lipids were altered in lamina propria neutrophils (Fig. 5D,E; Tables S12–S15). TSP1/TGF-β1 inhibition also reshaped lipid class composition, most prominently in epithelial neutrophils (Fig. 5F), leading to a pronounced increase in phospholipid classes, especially phosphatidylcholines (PC) and phosphatidylacids (PA) that contain an AA acyl chain and could thus serve as a metabolite precursor essential for LTB4 production.

**Figure 5.**
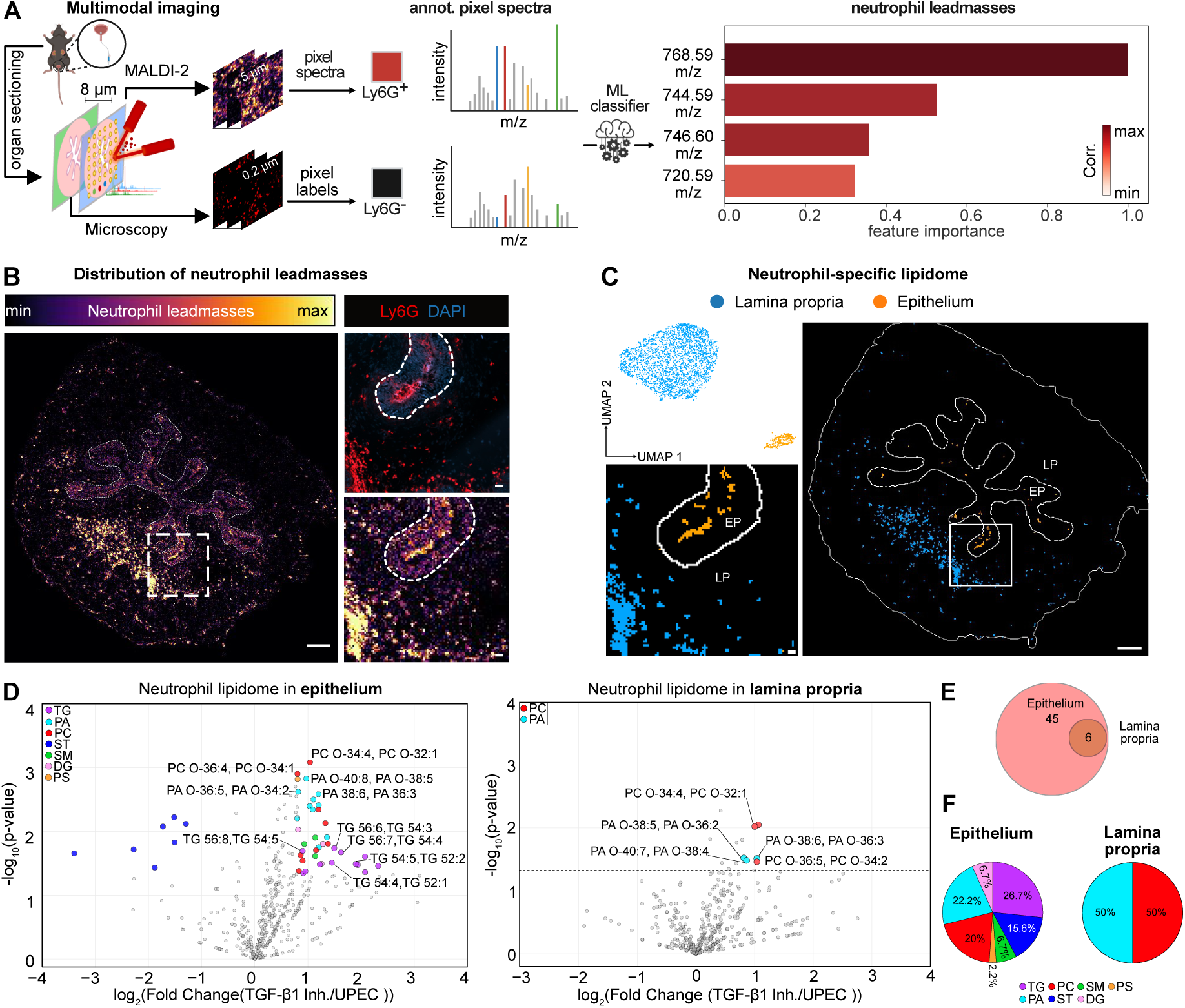
Spatial lipid analysis reveals TGF-β1–dependent neutrophil lipid signatures in the epithelium. **(A)** Schematic overview of the multimodal imaging workflow combining MALDI-2 mass spectrometry imaging and fluorescence microscopy. Organs are sectioned and imaged to acquire pixel-resolved mass spectra and immunofluorescence-based cell type labels (Ly6G for neutrophils). Machine learning (ML) classification is used to identify lipids associated with Ly6G⁺ neutrophils, revealing key lead masses based on feature importance. Partially created in BioRender. Widera, L. (2025) https://BioRender.com/t86f70g. **(B)** Concatenated spatial distribution of leadmasses identified in (A) and Ly6G^+^ neutrophils in consecutive tissue sections. Scalebar indicates 200µm (overview) and 20µm (zoom). **(C)** Dimensionality reduction of neutrophil-specific lipid signatures identifies two distinct clusters. Spatial localization indicates that one cluster aligns with epithelial neutrophils and the other with lamina propria neutrophils. Scalebars indicate 200µm (overview) and 20µm (zoom), respectively. **(D)** Differentially regulated lipid signatures in neutrophils in epithelium (left) and lamina propria (right). Lipids with log_2_(Fold Change)>0.8 and -log_10_(p-value)>1.3 and 2 are annotated. **(E)** Overlap of lipidomic signatures between neutrophils in epithelium and lamina propria. **(F)** Distribution of regulated lipid classes in neutrophils from each tissue compartment upon TSP1/TGF-β1 inhibition.

Next, we investigated the role of the AA metabolism in neutrophil cluster formation (Fig. 6A). To achieve this, we examined the distribution of the five most differentially regulated PC and PA phospholipids that by their *m/z* likely contain the AA metabolite precursor of LTB4 in their side chains. We observed that these phospholipids exhibited a strong correlation with neutrophil distribution in the epithelium (Fig. 6B). Since unsaturated fatty acid side chains of phospholipids contain the AA metabolite precursor for LTB4 production, we investigated the ability of neutrophils to convert the AA precursor into LTB4 through expression of Alox5 and the Alox5-activating protein (Alox5ap). To achieve this, we isolated neutrophils and surrounding epithelial cells from tissue sections using laser capture microdissection (LMD) (Fig. 6C) and analysed the expression of Alox5 and Alox5ap among other proteins by LC-MS/MS proteomics (Fig. 6D, Table S16). The analysis of epithelial regions revealed that neither epithelial cells close to neutrophil-rich regions nor distant epithelial cells expressed Alox5 and Alox5ap. However, neutrophils showed strong expression of both proteins (Fig. 6D), indicating that neutrophils harbour the enzymatic machinery required for LTB4 production within the specific epithelial tissue niche. To assess the role of LTB_4_ for neutrophil cluster formation and host defence, we pharmacologically inhibited AA-mediated LTB_4_ biosynthesis using Zileuton, a specific Alox5 inhibitor (Fig. 6E). Pharmacological inhibition of Alox5 significantly reduced neutrophil cluster formation within the epithelium (Fig. 6E, F). In addition, the reduced number of epithelial neutrophils was associated with an increased bacterial load in the bladder tissue (Fig. 6G). These findings highlight the critical role of TSP1/TGF-β1-driven lipid metabolism for neutrophil clustering and effective bacterial clearance.

**Figure 6.**
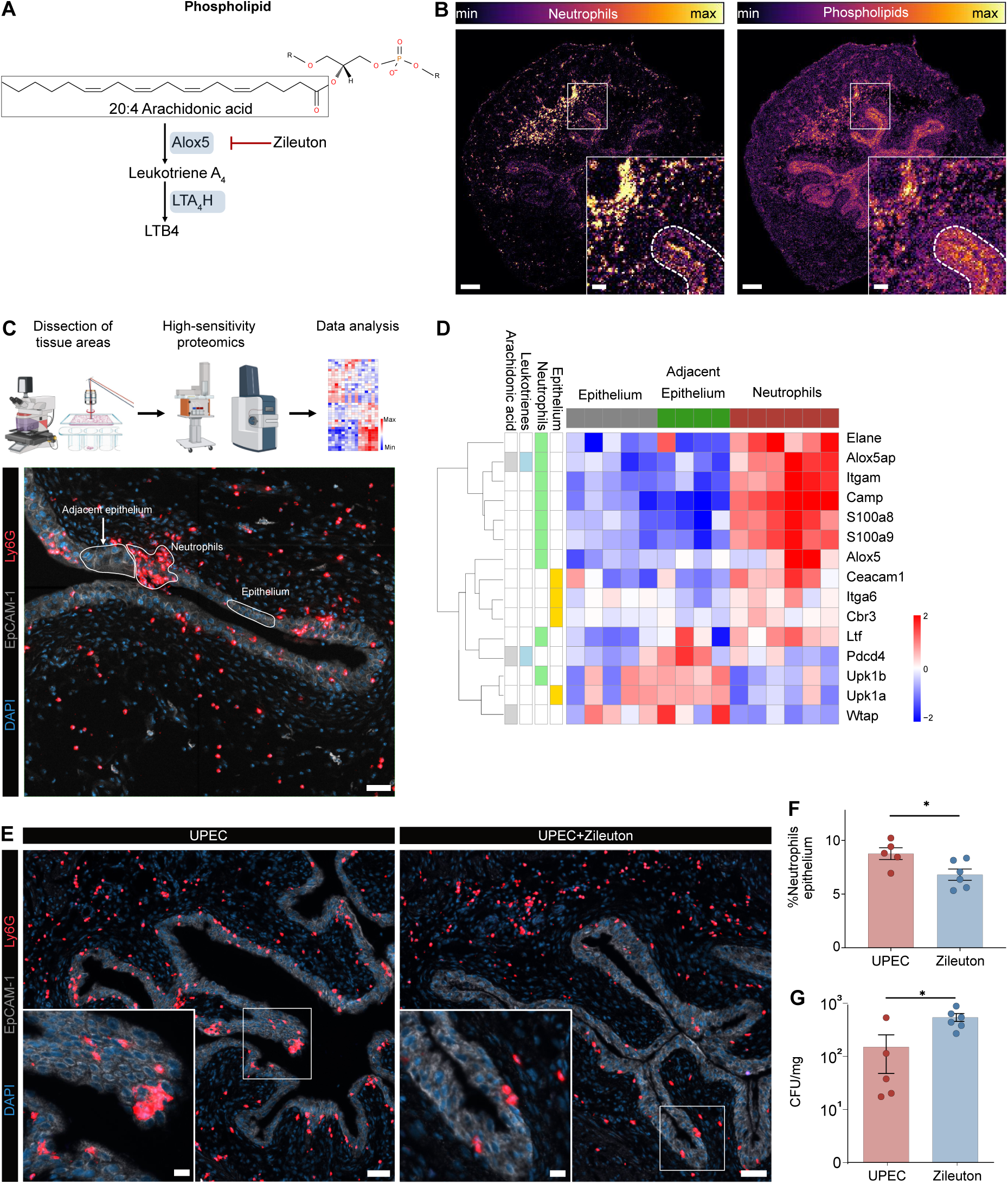
LTB₄ signaling promotes neutrophil clustering and bacterial clearance in UTI. **(A)** Structural illustration of AA-containing phospholipids and LTB_4_ synthesis. **(B)** MALDI-MSI images showing the spatial distribution of neutrophils (using the leadmasses identified in Fig. 5) and potential AA-containing phospholipids. Scalebars indicate 200µm (overview) and 50µm (zoom), respectively. **(C)** Schematic illustration of DVP, and representative image for areas used for LMD. Scalebar indicates 20µm. Partially created in BioRender. Widera, L. (2025) https://BioRender.com/nrdce7l. **(D)** LC-MS/MS proteomics of the distinct tissue areas were performed. Abundance distribution of proteins involved in AA metabolism, leukotrienes metabolism, neutrophils and epithelium, with p-values <0.05 (Kruskal-Wallis). For proteins ≥0.05, the p-value is shown. **(E)** Analysis of neutrophil distribution in histological sections of infected urinary bladders after inhibition of Alox5 using the pharmaceutical inhibitor Zileuton. Scalebars indicate 50µm (overview) and 20µm (zoom), respectively. **(F)** Quantitative results of neutrophil distribution within the epithelium. **(G)** Colony forming units (CFU) were determined using bladder homogenates. *p<0.05, Mann-Whitney; Data are mean±SEM (F,G).

## Discussion

Neutrophil swarming behaviour is a spatially confined multicellular response, characterized by coordinated neutrophil migration and cluster formation.^7,14^ This process is visible in many malignant situations, such as infections, and there is growing evidence for this process in other disease entities, such as tumour.^28^ In this study, we identify a spatially confined stroma-immune circuit in which TSP1-dependent activation of TGF-β1 licenses AA metabolism and neutrophil swarming.

A central feature of neutrophil swarm formation is its biphasic organization: initiation by pioneer neutrophils responding to danger-associated molecular patterns or early chemoattractants,^23^ followed by signal amplification through LTB₄-mediated recruitment of additional neutrophils.^7,14^ The importance of LTB₄ in this amplification step has been demonstrated in genetic ablation studies targeting either Alox5 and LTB₄ receptor, which resulted in defective swarming and pathogen control.^7,15,29^ Our findings extend these principles and uncover a novel upstream regulatory axis in which epithelial-derived TSP1/TGF-β1 signalling licenses LTB₄ biosynthesis in neutrophils in specific tissue niches. This positions epithelial cues not only as initiators of immune cell recruitment, but also as active modulators of the lipid signalling landscape.

Although neutrophils express TGF-β receptors and are highly responsive to this pleiotropic molecule,^30^ TSP1/TGF-β1 signalling may also be mediated through intermediate cell populations in the tissue microenvironment. One plausible scenario involves stromal fibroblasts of the lamina propria, which are well-known TGF-β1–responsive cells and could amplify epithelial signals by releasing lipid mediators or chemokines that promote neutrophil priming. Similarly, resident macrophages possess the machinery for arachidonic acid metabolism.^31,32^ Given their established role in chemokine signalling,^23^ macrophages may act as cellular relays that either directly produce leukotrienes or shape the inflammatory milieu to enable robust neutrophil swarming.

Our identification of TSP1 as a spatial organizer of neutrophil clustering highlights a previously unrecognized role for this matricellular protein in innate immunity, extending its established function as a regulator of latent TGF-β1 activation to the orchestration of neutrophil swarming. This spatial mechanisms mirrors prior findings that neutrophil swarm architecture is shaped by local tissue topography and microenvironmental constraints.^2,18^ The loss of epithelial-neutrophil circuits upon inhibition of the TSP1/TGF-β1 axis is associated with reduced neutrophil swarming and increased bacterial burden, underscoring the functional relevance of this spatially confined defence mechanism.

Multimodal imaging using spatial lipid profiling and fluorescence microscopy provided spatially resolved insight into the lipid-driven mechanism of neutrophil cluster formation. Specifically, accumulation of ether-linked PC species containing arachidonic acid upon TGF-β1 blockade indicated impaired use of AA toward LTB₄ synthesis. These observations align with mechanistic data from previous studies, showing that TGF-β1 modulates leukotriene biosynthesis at multiple levels: it increases the protein levels of cPLA_2_ and activates it via phosphorylation at Ser505, potentially through the mitogen-activated protein kinases p38 and ERK1/2, which are activated downstream of non-canonical TGF-β1 signalling.^33,34^ Beyond its effect on cPLA₂, TGF-β1 also enhances Alox5 activity through phosphorylation at Ser271 and Ser663, likely mediated by MAPK2 and ERK1/2.^35^ In addition, TGF-β1 promotes Alox5 transcription through a SMAD3/4-responsive element in its promoter, an effect further enhanced by 1,25-dihydroxycholecalciferol (vitamin D₃) and accompanied by upregulation of the essential cofactor FLAP (Alox5ap).^36–39^ In line with this, pharmacological inhibition of Alox5 resulted in defective neutrophil clustering and impaired bacterial clearance. Notably, this occurred without major changes in neutrophil extravasation into the stroma and overall maturation status, suggesting that lipid-mediated spatial organization, rather than developmental state, dictates neutrophil migration in this context.

Neutrophil swarming has emerged as a conserved multicellular behaviour observed across tissues and pathological contexts.^28,40^ Our study specifically delineates a TSP1/TGF-β1–dependent epithelial circuit that operates in the urinary epithelium during infection. Beyond infection, neutrophil clustering also emerges in cancer biology, where neutrophils either fight or support the tumour, and form heterotypic clusters with tumour cells influencing metastatic dissemination.^41^ Our identification of a tissue-derived checkpoint, in which TSP1-dependent activation of TGF-β1 licenses neutrophils, positions TGF-β1 not only as a determinant of tumour-associated neutrophil phenotype ^42^, but also as a spatially acting tissue signal that can tune neutrophil collective behaviour. By revealing how a matricellular regulator (TSP1) and TGF-β converge on neutrophil lipid metabolism to permit swarming, our data suggest a unifying principle whereby local tissue niches instruct conserved neutrophil programmes across infection and malignancy. Whether analogous epithelial–neutrophil communication pathways regulate swarming in other barrier tissues such as the intestine, skin, or tumour epithelia remains an open question and will require dedicated investigation.

Taken together, our findings define a previously unrecognized TSP1/TGF-β1–leukotriene axis that spatially organizes neutrophil swarming. This work expands the conceptual framework of neutrophil swarming by placing stroma-immune communication at the centre of lipid metabolism. This spatial coordinated coupling of epithelial signalling with neutrophil lipid metabolism opens new therapeutic avenues for modulation neutrophil accumulation and swarming.

## Methods

### Animal studies

Six- to eight-week-old female C57BL/6J mice were used throughout the experiments. Animals were purchased from Charles River Laboratories and maintained under specific pathogen-free conditions in the central animal facility at the University Hospital Essen. Mice were kept at 20–24 °C, 45–65% humidity in a 12 h dark-light cycle. The local review board (Bezirksregierung Köln, Landesamt für Verbraucherschutz und Ernährung in NRW / Recklinghausen, Germany) approved the animal experiments.

### Urinary tract infection model

UPEC strain 536 (O6:K15:H31) was cultured for 3 h at 37 °C in LB medium. The bacteria were harvested by centrifugation at 1500 × g for 20 min and the OD_600_ was measured. Bacteria were resuspended at a concentration of 10^10^ bacteria/ml in sterile PBS. A mixture of ketamine and xylazine 80 and 10 mg/kg body weight, respectively, dissolved in PBS was injected intraperitoneally to anesthetize the mice. The infection was performed by transurethral inoculation of 5 × 10^8^ UPEC in 0.05 mL PBS using a soft polyethylene catheter.

### TSP1/TGF-β1 inhibition *in vivo*

TSP1/TGF-β1 inhibitor was purchased from Hycultec and dissolved in sterile PBS at a concentration of 1.66 µg/ml and stored at - 80°C. For inoculation into the urinary bladder, the stock solution was diluted to a concentration of 0.6 µg/ml using sterile PBS. Mice were anaesthetized using a mixture of ketamine and xylazine (80/10 mg/kg body weight) by intraperitoneal administration. A soft polyethylene catheter was moistened with Instillagel and then carefully inserted through the urethra into the bladder. Subsequently, 50 µl of the inhibitor solution corresponding to 30 µg was inoculated 1 h post infection into the urinary bladder. The control group received the same volume of PBS.

### Inhibition of Alox5 *in vivo*

Zileuton was dissolved in a mixture of 10% DMSO, 40% PEG300, 5% Tween80 and 45% saline. Mice received two *i.p.* injections of Zileuton 23 h prior to UPEC infection and 1 h after infection. The Zileuton-treated mice received 35 mg/kg body weight, while the control mice received the same volume of the mixture of DMSO, PEG300, Tween80 and saline as indicated above.

### Determination of bacterial burden and TSP1 in tissue homogenates

Bladders were homogenized in 500 µl of sterile PBS, supplemented with protease inhibitor using a bead mill with 1.4 mm ceramic beads. To determine the bacterial burden, the homogenate was serially diluted in sterile PBS and plated on selective agar plates for 12 h at 37°C (ChromeID CPS Elite). To determine the amount of TSP1, an ELISA was performed according to manufactureŕs instruction. Amounts of TSP1 were normalized to overall protein amount in the homogenates using BCA-Protein Assay.

### Preparation of bladder single cell suspension for flow cytometry

Bladders were cut into small pieces with scissors and incubated at 37°C for 15 min with 100 mg/ml DNAse I and 0.5 mg/ml Collagenase Type I in RPMI 1640 medium supplemented with 10% heat-inactivated FCS with orbital rotation at 20 rpm. Tissue pieces were homogenized by vigorous pipetting, followed by a second incubation at 37°C for 15 min and rotation, and termination by the addition of 1 ml ice-cold PBS. Samples were homogenized by vigorous pipetting and filtered through a 70 µm cell strainer.

Single cell suspensions were incubated with antibodies in staining buffer (Cell Staining Buffer) for 20 min, 4°C, dark. The following antibodies were used for spectral flow cytometry at the indicated dilutions; CD101–APC (1:100), CD115–Alexa Fluor 488 (1:100), CD117–BV421 (1:50), CD11b–BUV496 (1:200), CD14–APC/Fire 750 (1:200), CD16–PE/Dazzle 594 (1:200), CD44–BV570 (1:300), CD45–PerCP (1:400), CD49d–BUV563 (1:100), CD62L–BV480 (1:200), CXCR2–PE (1:50), Ly6C–BV785 (1:300), Ly6G–BUV395 (1:300), MHC-II–BUV661 (1:100), CD170–BUV737 (1:100), and a viability dye—Zombie-UV (1:200). Incubation was stopped by adding 1 ml of PBS and the samples were centrifuged to collect the cells (400 g, 5 min, 4°C). Cells were fixed in 400 µl 4% PFA (10 min, RT, dark). Fixation was stopped by adding 1 ml PBS and the samples were centrifuged to collect the cells (400 g, 5 min, 4°C). Cells were resuspended in 150 µl PBS and measured using a Cytek Aurora full spectrum cytometer.

### Immunofluorescence microscopy

Bladders were fixed overnight in PLP buffer [pH 7.4, 0.05 M phosphate buffer containing 0.1 M L-lysine, 2 mg/mL sodium periodate and paraformaldehyde with a w/v concentration of 1%], equilibrated in 30% sucrose for 24 h and stored at −80 °C. Bladder tissue was cut into 8 µm thick sections at−20 °C using a cryostat. Unspecific binding was blocked by incubation of the sections with PBS containing 1% BSA and 0.05% Triton X-100 for 1 h. For antibody staining, the sections were heat-fixed at 70°C for 10 min, followed by rehydration and blockage of unspecific binding with blocking buffer (PBS+1%BSA+0.05% Triton X-100) (500 µl/slide, 1 hour, RT). Tissue sections were enclosed by a hydrophobic marking pen and incubated with antibodies against TSP1 (1:200), EpCAM-Biotin (1:200), Ly6G-AF647 (1:200) for 1 hour at RT in the dark. After washing (PBS+0.05% Triton X-100), sections were incubated with secondary antibodies Streptavidin-AF488 (1:400) and α-rabbit-AF568 (1:400) for 1hour at RT in the dark. After washing, nuclei were counterstained using DAPI (1:5000) for 5 min at RT in the dark. 2–3 sections per bladder were imaged on a Zeiss AxioScan.Z1.

An intensity threshold was used to generate masks for each fluorescent channel, cells were segmented using the StarDist plugin.^43^ The binary information for cellular and nuclear signals was coregistered. Automated analysis of cell densities was performed by a Java based algorithm. Using ImageJ, overlapping mask regions were used to identify cells, which were marked in the program with a point at the centre of the DAPI^+^ cell nucleus. The bladder tissue was segmented into lumen, epithelium, and lamina propria by employing the EpCAM-1 signal and cell densities were calculated.

For cyclic multiplex immunofluorescence microscopy through MACSima, bladder sections were mounted into MACSwell sample carriers, blocked using a blocking buffer containing 10% BSA and 2% goat serum for 1 at RT before nuclei were counterstained using DAPI-staining solution according to the manufacturer’s recommendations, and placed into a MACSima imaging system. Sections were then incubated with directly conjugated antibodies against Ly6G. Acquired fluorescence images were stitched using the pre-processing pipeline in MACS iQ View Analysis Software for downstream analysis.

### Cluster algorithm

Pairwise distances between cells were determined using a weighted depth first search to account for tissue boundaries. The distance matrix was transformed using a sigmoid function to emphasise close distances in form of a similarity matrix and pairs with values below 0.001 were treated as no similarity.

Clusters resulted from applying Markov Cluster algorithm of Clustering.jl (v0.15.7) with inflation of 1.2. The cluster area was calculated from the dilated boundaries of their alpha shapes generated with ɑ=80 using AlphaShapes.jl (v0.3.0). Significance of clusters was estimated using a hypergeometric distribution around the probability of seeing at least n cells in an area of size a given that a total of N cells are in the image with a valid area A, meaning tissue that is not more than 35 µm away from any cell.

### Light-sheet Microscopy

4 h prior sacrifice of mice, 10 µg of antibodies against EpCAM-1 AF488, Ly6G AF594 and CD31 AF647 were inoculated into the tail vein. Mice were euthanized using a ketamine/xylazine injection solution (200 mg/kg ketamine and 20 mg/kg xylazine) by intraperitoneal injection, followed by transcardial perfusion with 15 ml of cold PBS and 15 ml of cold 4% PFA solution. Following transcardial perfusion, mouse bladders were dissected and post-fixed in 4% PFA for an additional 4 h on ice. Samples were then dehydrated according to the ECi-clearing procedure (in a graded ethanol series with pH adjusted to 9,0 (50%, 70%, and two changes of 100%, each for 24 h) at RT.^44^ For optical tissue clearing, dehydrated specimens were immersed in ethyl cinnamate at RT until complete transparency was achieved. ECi-cleared bladders were imaged using a UltraMicroscope Blaze, equipped with a supercontinuum white light laser (460–800 nm) and an Andor Sona 4.2B-11 sCMOS camera (pixel size: 6,5 x 6,5 µm²). Samples were placed in ECi within a quartz imaging cuvette and imaged using defined excitation/emission filter combinations for the respective fluorophores: EpCAM-1 AF 488 (ex 500/20 nm; em 535/30 nm), Ly6G AF594 (ex 560/40 nm; em 620/60 nm), and CD31 AF647 (ex 630/60 nm; em 680/30 nm). Autofluorescence was recorded using ex 470/30 nm and em 525/50 nm. Overview scans were acquired using a 4x objective (NA 0,35) with 0,66x optical zoom and a z-step size of 4 µm. Higher-resolution image stacks were acquired with the same objective at 2,5x optical zoom and a z-step size of 3 µm. LSFM image stacks were imported into Imaris software (Bitplane, v9.7.1) for 3D reconstruction and quantitative analysis. A surface rendering of the EpCAM signal was generated to delineate the epithelium and optically isolate it from the underlying lamina propria of the bladder. The same surface rendering-based approach was used to segment Ly6G⁺ neutrophils and cell clusters within the optically isolated epithelium. Volumetric measurements of both epithelial compartment and neutrophil clusters were extracted and exported for each sample for downstream statistical analysis. To classify neutrophil clusters within the epithelium, volumetric data from Imaris-based surface renderings of single Ly6G⁺ cells and cell clusters were further analysed. First, the average volume of a single neutrophil was determined by selecting a field of view within the bladder devoid of visible clustering. In this region, individual neutrophils were segmented and rendered to provide a reference volume for a single cell. Based on this average neutrophil volume, cluster sizes were calculated by dividing the total surface volume of each segmented object by the single-cell reference value, thereby estimating the number of cells per cluster. Clusters were then categorized into three groups according to their estimated cell counts: small clusters (10 - 50 neutrophils), medium clusters (50 - 100 neutrophils) and large clusters (>100 neutrophils). Cluster count data were analysed using a Bayesian negative binomial generalized linear mixed model (NB-GLMM) implemented in Python using Bambi (v.0.15.0) and arviz (v.0.22.0). The model included fixed effects for experimental groups UPEC vs TGF-β1, size and their interaction, and a random intercept for each image to account for image-dependent effects. The model was fit with 2000 posterior draws, 4000 tuning steps, 4 computational cores and random_seed=9, using weakly informative priors for all parameters. Posterior distributions were summarized by mean, standard deviation, and 95% highest density intervals (HDI), and posterior probabilities of positive or negative effects were computed. For further interpretability, approximate z-scores and one-sided p-values were calculated from posterior means and standard deviations. Additionally, the posterior draws were exponentiated to obtain values on the natural scale, and the corresponding mean, standard deviation, and 95% highest density intervals were computed.

### Fluorescence *in situ* hybridization

Bladders were snap frozen in 2-methylbutan on dry ice using a spatula for 5 s and then stored at −80°C for further analysis. RNA probes for TSP1 (Thbs1, NM_011580.4) and TGF-β1 (TGFB1, NM_011577.2), were purchased from Molecular Instruments and staining was performed according to manufactureŕs instructions. Sections were imaged at Zeiss Axio-Observer.Z1.

### Sample preparation and acquisition of MALDI-MSI data

The workflow for the acquisition, analysis and co-registration of MALDI data and microscopy images, as well as MALDI-MSI data, was recently established by the msiFlow pipeline.^45^ Briefly, bladders were isolated 24 h after infection with UPEC and snap-frozen in 2-methylbutan and sectioned to 8 µm thickness using a cryostat. Sections were thaw-mounted on SuperFrost™ slides, vacuum-sealed, and stored at –80 °C (or on dry ice in case of transport) until further use. For MALDI, they were under a stream of dry N_2_ gas, and 2,5-dihydroxyacetophenone (DHAP) matrix was applied onto the tissue section by sublimation using a home-built sublimation device described earlier.^46^ Samples were transferred to the MALDI ion source immediately after matrix application. MALDI MSI measurements were performed with a timsTOF fleX MALDI-2 instrument with microGRID extension. The positive-ion mode was used, and all reported data acquired with a pixel size of 5 µm x 5 µm. MALDI and post-ionisation lasers (MALDI-2) were operated with a pulse repetition rate of 1 kHz and an inter-laser delay of 10 µs. For MALDI-2 measurement, the ablation laser power was set to 80% (instrument readings) and 25 shots were set per pixel to acquire full mass spectra in the *m/z* range of 300 to 1500. All standard MALDI-2-MSI measurements were performed with the trapped ion mobility spectroscopy (TIMS) option of the instrument being disabled. MS/MS measurement, facilitated by TIMS separation, was performed using a 50 µm pixel size and 250 laser shots. The 1/k_0_ range was from 1.4 to 1.8 and nitrogen (N_2_) was used as the collision gas. The ramp time was set to 250 ms, the isolation window to 1 Da and the collision energy to 30 eV. MALDI-DDA-MSI was used to validate tentatively identified lipids by on-tissue MS/MS analysis by alternating between full scan and MS/MS pixels.^47^ Front illumination was used with a step size of 10 µm in the x-direction and 20 µm in the y-direction. For full scan pixels the ion detection range was set to m/z 550-1500. For DDA MS/MS measurements, an isolation window of 1 Da was used with a fixed first mass of m/z 100 and a normalized collision energy of 25. The exclusion time was set to 30 s with isotope exclusion enabled. The mass resolution for both full scan and MS/MS was set to 70,000 and the injection time was fixed at 250 ms. MALDI-DDA-MSI data were processed using Lipostar MSI (version 1.3) and annotated based on the Lipid Maps Structure Database.

### Analysis of MALDI-MSI data

MALDI-MSI data were pre-processed using custom Python scripts automated via a Snakemake pipeline, which processed raw timsTOF files in parallel and generated imzML files along with quality control visualizations. Savitzky-Golay smoothing was applied to reduce spectral noise, and centroid spectra were extracted using SciPy’s find peaks function, selecting peaks with an SNR of at least three. To correct for possible mass drifts during the MSI run, a kernel-based alignment method (adapted from pyBASIS) was used to align peaks present in at least 3% of all pixels. Off-tissue matrix pixels were detected using UMAP and HDBSCAN clustering, and subsequently removed based on boundary connectivity and Spearman correlation (>0.7). A binary image refinement step removed small artefacts before final matrix removal. Spatial coherence was calculated to assess ion information, with low coherence ions removed. Data were then normalized using median-fold change normalization to correct for intra- and inter-sample variation. Outlier detection was performed using a UMAP-HDBSCAN-based approach, identifying sample-specific-clusters where most pixels were from a single sample. Finally, de-isotoping was performed iteratively, matching isotopes based on m/z tolerance and theoretical intensity patterns. This workflow was in part described recently. ^45^

### Lipid annotation

Initial tentative molecular species annotations were generated using the bulk structure search on the LipidMaps website, matching various lipid classes, such as free fatty acids (FA), ceramides (Cer), sphingomyelins (SM), Hexosylceramids (HexCer), triacylglyerols (TG), diacylglycerols (DG), PC, PA, phosphatidylserines (PS), phosphatidylethanolamines (PE), phosphatidylglycerols (PG), sulfatides (ST) with precursor ions [M + H]+, [M-H_2_O + H]+, [M + Na]+ and [M + K]+ within a ±0.01 *m/z* tolerance. The list was then manually refined based on biological likelihood.

### Lipid profiling of neutrophils

Neutrophil-specific leadmasses were determined by combining fluorescence microscopy and MALDI-MSI. To this end, consecutive tissue sections were measured by MALDI-MSI and immunofluorescence microscopy (Ly6G staining for neutrophils). Registration of MALDI-MSI and microscopy data and a machine learning classifier was used to identify molecular species (mass-to-charge (*m/z*) values) that showed a strong spatial colocalisation with Ly6G^+^ neutrophils. These leadmasses served as a proxy for neutrophil distribution in the MALDI-MSI dataset and were used for further neutrophil-related lipid profiling. The combined leadmass images were generated using the mean of the 4 identified lipid intensities, followed by contrast enhancement using percentile stretching. Next, the generated leadmass images were segmented by Yen thresholding generating a binary image. Small artefact regions < 3 pixels were removed from these thresholded images. To distinguish between neutrophil pixels in the epithelium and the lamina propria/muscle tissue, unsupervised tissue segmentation of the MALDI MSI data was performed by spatial k-means implemented in msiFlow.^45^ The lipidomic profile of neutrophil pixels in the epithelium and the lamina propria/muscle tissue between UPEC+TGF-β1 inhibitor and UPEC-infected mice were statistically analysed using msiFlow. Therefore, the median spectra of the pixels were extracted and statistically analysed by two-sided statistical tests using the Python package SciPy. The standard *t*-test was used for normally distributed populations with equal variances and the Welch’s *t*-test for normally distributed populations with unequal variances. Non-normally distributed populations were statistically analysed by the Wilcoxon rank-sum test. The Shapiro–Wilk test and Levene test were used to test for normal distribution and equal variances. The ratio between the means of two populations (here UPEC+ TGF-β1 inhibitor and UPEC) was determined as log_2_(fold change).

### Sample preparation for LC-MS/MS proteomics

For bulk bladder proteomics, a halved bladder frozen in 2-methylbutane was homogenized using a bead mill (1.4 mm ceramic beads) in 100 µl ddH_2_O supplemented with Complete mini-EDTA free protease inhibitor and adjusted with 100 µl double-concentrated lysis buffer to a final concentration of 50 mM Tris-HCl, pH 7.8, 150 mM NaCl, and 1% SDC. The samples were sonicated (2 × 5 min on wet-ice) and centrifuged (16000 g, 5 min, 4°C). The protein concentration of supernatant was determined using BCA assay according to the manufacturer’s instructions. For tryptic digestion, 20 µg of proteins were reduced with 10 mM DTT at 56 °C for 30 min, followed by cysteine alkylation with 2-iodoacetamide (final concentration 20 mM) for 30 min at RT in the dark. Tryptic digestion was performed, using the single-pot, solid-phase-enhanced sample preparation (SP3) protocol, as previously described.^48^ In brief, 20 µg of protein was bound to hydrophilic and hydrophobic beads (50:50 (w/w); using a bead to protein ratio of 10:1 for 18 min at RT. Samples were washed twice with 100% acetonitrile (ACN), twice with 70% ethanol and then reconstituted in 50 mM ammonium bicarbonate buffer. Tryptic digestion was performed at a trypsin-to-protein ratio of 1:100 for 16 h at 37 °C. Peptides were eluted twice with 20 µl 1% formic acid, 2% dimethyl sulfoxide in water. Peptides were vacuum-dried in a Concentrator plus (Eppendorf SE, Hamburg, Germany) and stored at −80 °C until further use. Directly prior to measurement, peptides were reconstituted in 0.1% of formic acid. The peptide concentration was estimated using a NanoDrop Microvolume Spectrometer.

For Deep Visual Proteomics (DVP), fresh-frozen bladder tissue was cryo-sectioned at 10 µm onto PPS metal-frame slides. Sections were immediately vacuum-dried and stored at −80°C for less than 3 days. For staining, slides were incubated with Ly6G-AF647 (1:200 in Intercept Blocking Buffer), washed three times with PBS, counterstained with DAPI (1:5000 in in Intercept Blocking Buffer), washed again three times with PBS, and rinsed with LC-MS–grade water. A total of 0.1 mm² of the region of interest was micro-dissected using a Leica 7 LMD instrument and collected into 0.5 ml protein low-binding tubes. Samples were centrifuged for 5 min at 16,000 × g and lysed in 10 µl of reducing lysis buffer (0.1% DDM, 5 mM DTT, 50 mM ammonium bicarbonate). Lysates were sonicated in an ice-water bath (2 × 5 min), briefly spun down (16,000 × g, 15 s), and reduced for 60 min at 60°C, followed by cysteine alkylation with iodoacetamide (final concentration 20 mM) for 30 min at 37°C in the dark.

Trypsin (0.3 µL of 10 ng/µl in 50 mM ABC) was added, and samples were incubated for 16 h at 37°C. Digestion was quenched with 15 µl of 10% formic acid. Peptides were dried in a Concentrator Plus and stored at −80°C. Immediately prior to LC-MS/MS analysis, peptides were reconstituted in 20 µl of 0.1% FA.

### LC-MS/MS parameters for proteome data acquisition

For bulk bladder proteomics, 300 ng of peptides, and for DVP proteomics 5 µl peptide solution in 0.1% formic acid were loaded onto an Evotip (single-use C18 StageTip-based trap columns designed for low-volume peptide cleanup) following the manufacturer’s instructions. Briefly, Evotips were activated with 0.1% formic acid in ACN, conditioned with 2-propanol, equilibrated with 0.1% formic acid, and then loaded for 1 min using centrifugal force at 800 g. Evotips were subsequently washed with 0.1% formic acid. Afterwards,100 µL of 0.1% formic acid was added to each tip to prevent drying.

Peptide separation was performed on an Evosep One system using the predefined 30SPD method, on an Aurora Elite CSI C18 UHPLC column (15 cm × 75 µm, i.d., 1.7 µm particle size, 120 Å pore size). Eluting peptides were injected into a TimsTOF^HT^ mass spectrometer, via a CaptiveSpray source held at 1600 V. The mass spectrometer was operated in data-independent acquisition (DIA) PASEF mode. The accumulation and ramp time for the dual TIMS analyser were set to 100 ms at a ramp rate of 9.42 Hz. MS data were acquired over an m/z range of 100–1700 and a TIMS ion mobility range of 1/K0 = 0.6–1.6. For DIA–PASEF, a standard long-gradient method with a cycle time of 1.8 s was used. For Data-Independent Acquisition-Parallel Accumulation–Serial Fragmentation (DIA-PASEF), a standard “Long gradient” method with a cycle time of 1.8 s was used. Within each cycle, a precursor mass range of 400 to 1201 Da and a mobility range of 0.6 to 1.6 1 K0^-1^ was fragmented. At MS1 level, one ramp was executed per cycle. For MS/MS measurements 32 MS/MS windows with a mass width of 26.0 Da each and a mass overlap of 1.0 were distributed across the 16 MS/MS ramps.

### Data extraction and analysis for proteome data

LC-MS raw data were processed using the directDIA algorithm in Spectronaut (Version 19.1.240806.626), applying BSG factory settings (Enzyme/ Cleavage rules: Trypsin/P; Fixed modification: Carbamidomethyl (C); Variable modifications: Acetyl (Protein N-term) and Oxidation (M)), against a reviewed murine UniProt/Swissprot FASTA database (downloaded on 29th of October 2024, containing 17221 target sequences). Peptides and proteins were considered identified with an FDR < 0.01 at peptide and protein level, respectively. For Deep visual proteomics (DVP) data missing values were imputed using the missing not at random (MNAR) hypothesis. Missing values were imputed from the normal distribution per protein, with a downshift of 1.3, toggling values across a range of 0.3 for each animal separately. Differentially abundant proteins between groups were determined through Kruskal-Wallis-test, accepting proteins with a p-value < 0.05 as significant across the tissue areas. Quantification was performed at the MS2 level, with imputation disabled and cross-run normalization enabled. Normalized protein abundances were exported and analysed in the R software environment (Version 4.4.2, R Core Team (2021) (Table S1). R: A language and environment for statistical computing. R Foundation for Statistical Computing, Vienna, Austria. Differentially abundant proteins between pairs of predefined phenotypes were determined through Student’s t-testing. Proteins, identified with a q-value (Permutation-based FDR) < 0.05 and at least 1.5-fold change difference between groups, were considered significant and subjected to further analysis. The standard *t*-test (t.test() function from the R “stats” package) was used for normal distributed data and the ratio between the means of two groups was determined as log2(fold change). Significantly regulated proteins (log_2_(fold change) ≥ ±0.5; p-value < 0.05) were further analysed using the STRING database (version 12.0). PCA was performed using scikit-learn (v.1.7.2). The full STRING network was used with all available active interaction sources (text mining, experiments, databases, co-expression, gene fusion, neighborhood, and co-occurrence). Only high-confidence interactions (score ≥ 0.700) were included. To identify functional protein modules, DBSCAN clustering was performed directly within STRING using default parameters. STRING enrichment analysis (Biological Process, GO) was performed for each cluster, with terms grouped by similarity ≥ 0.8 and filtered at FDR ≤ 0.05. Top 10 terms were displayed, sorted by signal. Heatmaps were generated using pheatmap^49^ and ggplot2^50^.

### NIRL-based 3D proteomics

Laser ablation was performed using an optical setup as previously reported.^51^. In brief, a nanosecond infrared laser (NIRL) system (Opolette SE 2731, Opotek, Carlsbad, CA, USA) was tuned to the wavelength of 2940 nm for cold tissue vaporization with single laser pulses of 7 ns pulse width. First, the laser beam is collimated and broadened by a 1:2 telescope, then focused by a 150 mm lens leading to a focus spot diameter of 100 µm at sample position, applying 730 µJ pulse energy. Between lens and sample, a dichroic mirror was introduced to integrate a monitoring camera path for aiming purposes. Layer-wise tissue ablation was performed utilizing a motorized stage (composed of MLT25, SPX-RLD4, XPS-DRV11), which was programmed to move in a meander pattern while triggering the laser with 20 Hz. The applied ablation pattern consists of 21 x 21 single laser shots, forming a layer with the dimensions of 2200 x 2200 x 23 µm³, which was determined using optical coherence tomography (OCT). We applied a layer-by-layer laser ablation starting from the inner side of the flattened formalin-fixed halved bladder. Each sampled layer was collected on in a separated well of a PTFE-coated objective glass slide, which was placed 1 mm above the sample. After ablating through the bladder, the condensed aerosol samples were processed for downstream proteomic analysis. The condensed aerosol was resuspended in 10 µl of 0.01% DDM in 25 mM ammonium bicarbonate. PTFE-coated slide was placed on a heating block for 5 min at 95°C, with LC-MS grade water added during the heating to maintain the samples in solution. Proteins were digested by adding 20 ng of trypsin and incubating overnight at 37°C in humidity chamber. Finally, the sample slides were dried in a vacuum centrifuge.

Chromatographic separation of peptides was achieved with a two-buffer system (buffer A: 0.1% FA in H_2_O, buffer B: 0.1% FA in 80% ACN) on a UHPLC (VanquishTM neo UHPLC system). Attached to the UHPLC was a PepMap™ Neo trap cartridge (300 µm x 5 mm, 100 Å pore size, 5 µm particle size, C18) for online desalting and purification, followed by a C18 reversed-phase column (75 µm x 250 mm, 130 Å pore size, 1.7 µm particle size, peptide BEH C18, nanoEase). Peptides were separated using a 55 min method with linearly increasing ACN concentration from 2.5% to 37.5% buffer B over 45 min. Eluting peptides were ionized using a nano-electrospray ionization source (nano-ESI) with a spray voltage of 1,800V and analysed on a quadrupole-orbitrap hybrid mass spectrometer (Exploris 480, Thermo Fisher Scientific). For the acquisition of the full scan spectrum, the maximum injection time was set to 240 ms or until a charge density of 3 x 10^6^ ions (AGC Target) was reached.

Fourier transformation-based mass analysis of the electronic data from the orbitrap mass analyser was performed covering a mass range of m/z 400 – 1200 with a mass resolution of 120,000 (at m/z 200). Within a precursor mass range of m/z 380 – 980, fragmentation in DIA-mode with 12 Da-wide isolation windows and a window overlap of 1 Da was performed at a normalized collision energy of 28% using higher energy collisional dissociation (HCD). The maximum injection time was set to 54 ms or until a charge density of 2 x 10^7^ ions (AGC Target) was reached. In these experiments, the orbitrap resolution was set to 30,000 and the scan range was set to m/z 350-2000.

LC-MS raw data were processed using the directDIA algorithm in Spectronaut (Version 19.1.240806.626), applying BSG factory settings (Enzyme/Cleavage rules: Trypsin/P; Fixed modification: Carbamidomethyl (C); Variable modifications: Acetyl (Protein N-term) and Oxidation (M)), against a reviewed murine UniProt/Swissprot FASTA database (downloaded on 29th of October 2024, containing 17221 target sequences). Quantification was performed at the MS2 level. Cross-run normalization and imputation were disabled. Unnormalized protein abundances were used for further downstream analysis, exported, and analysed in the R software environment (v 4.1.2; R Core Team 2021).

Protein abundances were log₂ transformed to approach the Gaussian probability distribution. Data normalization to compensate for injection amount discrepancies was achieved by subtracting the mean abundance per sample for each protein quantified in the respective sample. Imputation was performed under the missing not at random (MNAR) hypothesis. Missing values were imputed from the normal distribution per protein, with a downshift of 1.3, toggling values across a range of 0.3 for each animal separately. Protein abundance-derived similarity clusters across laser ablation layers were determined by Monte Carlo reference-based consensus clustering (M3C)^52^ for each animal. The maximum number of clusters was set to 12. Hierarchical clustering was used as a clustering method for 25 iterations, sampling 80% of all samples per iteration. Reverse PCA and Cholesky decomposition–based resampling was used, respectively, for the generation of a null reference distribution for 100 Monte Carlo simulations. For the real data set, 100 simulations were applied. Cluster stability was determined by entropy. Principal components (PCs) of ablation layers were created using a nonlinear iterative squared principal component analysis (NIPALS-PCA) with a maximum of three components. M3C clusters were assigned to cellular layers by analysing the scaled (0 to 1) protein abundance per animal to established marker proteins for different bladder cell layers.^27^ A score for each cellular layer was calculated by averaging the scaled abundance for each marker, associated with a respective cellular profile for M3C cluster and animal. If a score > 0.9 was reached, a cell type was considered identified. The Pearson correlation of TSP1 to all identified proteins and specifically known TSP1-interactors^53^ was calculated based on log2 transformed, normalized and imputed data separately for UPEC infected and healthy bladders. Proteins, positively correlating with a strong Pearson correlation coefficient >70% and a p-value < 0.001 were considered significant. To visualize known associations between corelating proteins to TSP1 in UPEC and/or healthy bladders, a STRING protein-protein interaction network was generated. Interactions between were accepted at a high confidence > 0.8. All interaction sources were enabled. To identify biological processes, statistically overrepresented in proteins correlating with TSP1 in UPEC and/or healthy bladders, enrichment analysis based on the gene ontology biological processes database (GO-BP)^54^ was performed, using the STRING app in the Cytoscape (Version 3.10.2^55^) environment. Terms identified as enriched with a q-value (Benjamini Hochberg FDR) < 0.05 were considered significantly enriched. Targeting the abundance distribution of selected gene sets across M3C clusters, the following data sources were used: Neutrophil Markers: Human Protein Atlas, single cell immune cell resource^56^; Arachidonic Acid metabolism: Kyoto encyclopedia of genes and genomes (KEGG^57^). Leukotriene metabolism: GO-BP. Within the R software environment, the following packages were used: *openxlsx*^58^, dplyr^59^, *gridExtra*^60^, *ggplot2*^50^, *tidyr*^61^, *mixOmics*^62^*, naniar*^63^*, lubridate*^64^*, M3C*^52^*, pheatmap*^49^, *RColorBrewer*^65^, *purrr*^66^*, stringr*^67^.

## Key resource table

| REAGENT or RESOURCE | SOURCE | IDENTIFIER |
| --- | --- | --- |
| **Antibodies and antibody binding agents** |  |  |
| CD101 APC | ThermoFisher Scientific, #12-1011-82, | RRID: AB_1210728 |
| CD115 AF 488 | Biolegend, #135502 | RRID: AB_1937293 |
| CD117 BV 421 | Biolegend, #105804 | RRID: AB_313213 |
| CD11b BUV496 | BD Bioscience, #569708 | RRID: AB_3685251 |
| CD14 APC/Fire 750 | Biolegend, #123332 | RRID: AB_2800591 |
| CD16 PE/Dazzle 594 | Biolegend, #158012 | RRID: AB_2876543 |
| CD44 BV 570 | Biolegend, #103037 | RRID: AB_10900641 |
| CD45 PerCP | Biolegend, #103130 | RRID: AB_893339 |
| CD49d BUV 563 | BD Bioscience, #741243 | RRID: AB_2870794 |
| CD62L BV 480 | BD Bioscience, #746726 | RRID: AB_2743990 |
| CD182 PE | Biolegend, #149304 | RRID: AB_2565692 |
| CD31 AF 647 | Biolegend, #623554 | RRID: AB_2161029 |
| Ly6C BV 785 | Biolegend, #128041 | RRID: AB_2565852 |
| Ly6G BUV 395 | BD Bioscience, #563978 | RRID: AB_2716852 |
| Ly6G AF 594 | Biolegend, #127636 | RRID: AB_2563207 |
| Ly6G AF 647 | Biolegend, #127610 | RRID: AB_1134162 |
| Ly6G PE | Miltenyi Biotec, #130-102-934 | RRID: AB_2660050 |
| MHC II BUV 661 | BD Bioscience, #750280 | RRID: AB_2874471 |
| CD170 BUV 737 | ThermoFisher Scientific #367-1702-82 | RRID: AB_2572866 |
| Zombie-UV Fixable Viability Kit | Biolegend, #423108 | RRID: AB_3676450 |
| a-thrombospondin-1 | Abcam, #ab267388 | / |
| CD326 Biotin | Biolegend, #118204 | RRID: AB_1134178 |
| CD326 AF 488 | Biolegend, #118210 | RRID: AB_3674977 |
| Streptavidin AF 488 | Invitrogen, #S11223 | RRID: AB_2315383 |
| Donkey a-rabbit | Invitrogen, #A10042 | RRID: AB_2534017 |
| Cell Staining Buffer | Biolegend | #420201 |
| TSP1 (Thbs1, NM_011580.4) Fish-probe | Molecular Instruments |  |
| TGF-β1 (TGFB1, NM_011577.2) Fish probe | Molecular Instruments | - |
| Mouse Thrombospondin 1 ELISA Kit | Hycultec | # AB-RK03230 |
| **Reagents and Instruments** |  |  |
| 0.5 ml screw cap tubes pre-filled with 1,4 mm ceramic beads | VWR International | #432-0375 |
| 2-Methylbutan | Sigma-Aldrich | Cat #320404-1L |
| 2-propanol | Sigma-Aldrich | #1027812500 |
| 2,5- dihydroxyacetophenone (DHAP) MALDI matrix | Merck | Cat #490-78-8 |
| Acetonitrile LC-MS Grade | Thermo Fisher Scientific | #51101 |
| Ammonium Bicarbonate | Sigma-Aldrich | #09830 |
| Aurora Elite CSI C18 UHPLC column | IonOpticks |  |
| Axio Scan Z.1 | Zeiss | - |
| Bead Mill Max | VWR International | - |
| Bovine serum albumin | GE Healthcare | Cat #K45-001 |
| C18 reversed-phase column | Waters |  |
| ChromID CPS Elite | Biomérieux | #43821 |
| Collagenase Type I | Sigma-Aldrich | #C2674-1G |
| Concentrator Plus | Eppendorf SE |  |
| DAPI | Life Technologies | Cat #D1306 |
| Dimethylsulfoxid (DMSO) | PanReac AppliChem | #A3672.0250 |
| Dimethylsulfoxid (DMSO) | Sigma-Aldrich | #D5879 |
| Dithiothreitol (DTT) | Roche | Cat #10708984001 |
| DNase I | Sigma-Aldrich | #D4513-1VL |
| mini-EDTA protease inhibitor | Sigma-Aldrich | #11836170001 |
| Ethanol, Ultra LC/MS | Carl Roth | #1HP8.1 |
| Ethylcinnamat (ECI) | Sigma-Aldrich | #112372-100G |
| Evosep One | Evosep | - |
| Evotip Pure, 10x96 tips | Evosep | #EV2013 |
| FBS Premium, South America origin, | PAN-Biotech | #P30-3302 |
| Formic acid | Biosolve | Cat #00069141 |
| Iodoacetamide (IAA) | Sigma | Cat #I1149 |
| L-lysine | Sigma-Aldrich | Cat #L5626 |
| LMD7 Laser Microdissection<br>Microscopes | Leica |  |
| Intercept (PBS) Blocking Buffer | LI-COR Bioscience | Cat #927-70001 |
| LSKL (TGF-β1 inhibitory peptide) | Hycultec | #HY-P0299 |
| MACSima | Miltenyi Biotec |  |
| NanoDrop Microvolume Spectrometer | Thermo Fisher Scientific |  |
| Orbitrap | Thermo Fisher Scientific |  |
| Paraformaldehyde | Sigma-Aldrich | Cat #P6148 |
| PepMap™ Neo trap cartridge | Thermo Fisher Scientific |  |
| PBS | Life Technologies | Cat #18912-014 |
| PPS metal-frame slides | Leica |  |
| Sequencing Grade<br>Modified Trypsin (5x20μg, lyophilized) | Promega | #V5111 |
| Sera-Mag™ magnetic carboxylate<br>modified particles (Hydrophobic) | Sigma-Aldrich | #GE44152105050250 |
| Sera-Mag™ Magnetic carboxylate<br>modified particles (Hydrophilic) | Sigma-Aldrich | #GE24152105050250 |
| Sodium Deoxycholate Detergent | Thermo Fisher Scientific | #89904 |
| TimsTOF Flex-MALDI2 | Bruker | - |
| Vanquish™ neo UHPLC system | Thermo Fisher Scientific |  |
| UltraMicroscope Blaze | LaVision BioTec |  |
| Experimental Models: Mouse Strains |  |  |
| C57BL/6 J | Jackson Laboratories | RRID:IMSR_JAX:000664 |
| Software |  |  |
| Affinity Designer | Serif |  |
| Fiji | ImageJ |  |
| FlowJo 10 | FlowJo |  |
| GraphPad Prism, version 10 | GraphPad Software |  |
| Lipostar MSI version 1.3 | Molecular Horizon |  |
| Python 3.12 | https://www.python.org/ |  |
| Spectronaut<br>(version 19.1.240806.626) | Biognosys |  |
| MACSiQ | Miltenyi Biotec |  |
| ZEN Digital Imaging for Light<br>Microscopy, ZEN 2012 | Zeiss |  |
| Imaris | Bitplane<br>(Oxford Instruments) |  |

## Data availability

The mass spectrometry proteomics data have been deposited to the ProteomeXchange Consortium via the PRIDE partner repository^68^. Data and supplementary tables (S1-S16) will be publicly available after the peer-review process.

## Author contributions

L.W., A.N., S.B., M.R., J.H., B.Si. and D.H., performed experiments

A.P., M.M., A.Go., S.T., S.S.T. and J.D. contributed to methodology and technical support.

L.W., A.N., P.S., D.S., D.H. and H.V. analysed data.

L.W., P.S., J.D., D.S., D.H., and H.V. prepared figures and visualizations.

J.So., K.D., T.C., J.J., H.S., O.So., A.G., T.L., O.S., M.G. and J.H. provided resources, reagents, and expertise.

L.W. and D.R.E. wrote the original draft.

D.R.E. supervised the work, acquired funding, provided project administration, and conceived and designed the study.

All authors contributed to review and editing of the manuscript.

## Acknowledgements

We acknowledge support by the Open Access Publication Fund of the University of Duisburg-Essen, the Central Animal Facilities of the Medical Faculty Essen, the Imaging Center Essen and the Immunoproteomics group, supported by INST 20876/486-1. The graphical abstract was created with support by Nina Knubel, Media Communication, University of Münster. We received funding from the German Research Foundation: FOR5427 SP1-SH542-3/1, ERA-NET NEURON (01EW2503) (OS); FOR5427 (466687329) SP4 (DRE); EN984/15-1, 16-1 and 18-1 (539301313) (DRE); TR296 (424957847) P09 (DRE); TR332 (449437943) A3 and Z1 (DRE), C5 (AG), A2 (OSo), A5 (JJ), B4 (TC), B6 (TL), C6 (MG), Z1 (OSo and KD); INST 20876/486-1 (DRE), 516868494 (HS), 518551069 (HS), 247354600 (HS), 247377969 (HS), 426788273 (HS) and the Mildred Scheel Cancer Carrer Center Hamburg (JH). MALDI-MSI measurements were supported by Bruker Daltonics.

## Supplementary Figures

**Figure S1.**
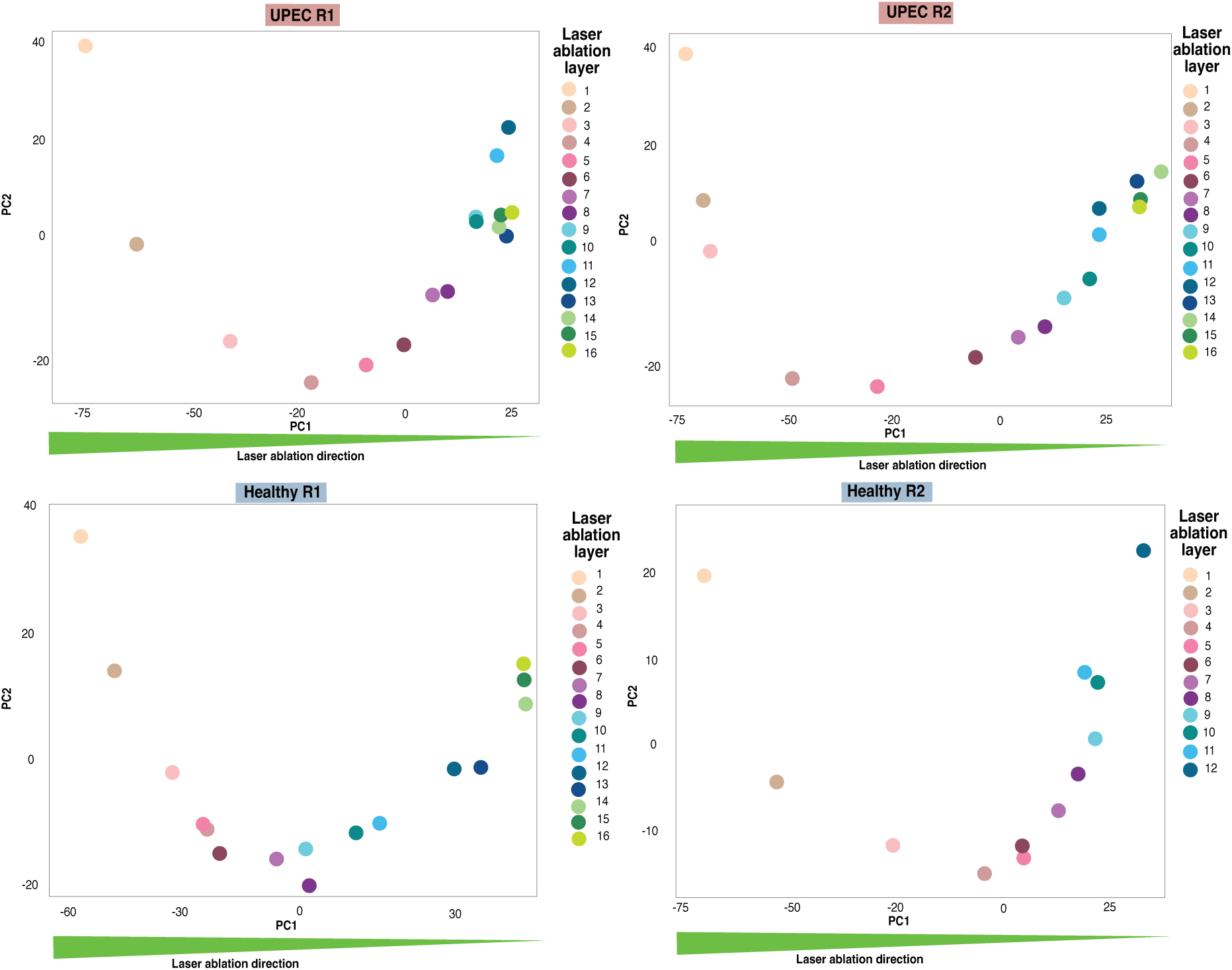
Unsupervised PCA of the consecutive laser ablation layers in NIRL 3D proteomics data. Scatter plot visualisation of the first two principal components (PCs) from unsupervised Principal Component Analysis (PCA), based on all quantified proteins per animal (Healthy R1: N =4036; Healthy R2: N=4078; UPEC R1: N=4010; UPEC R1: N=4085). Colours represent the different consecutive laser ablation layers.

**Figure S2.**
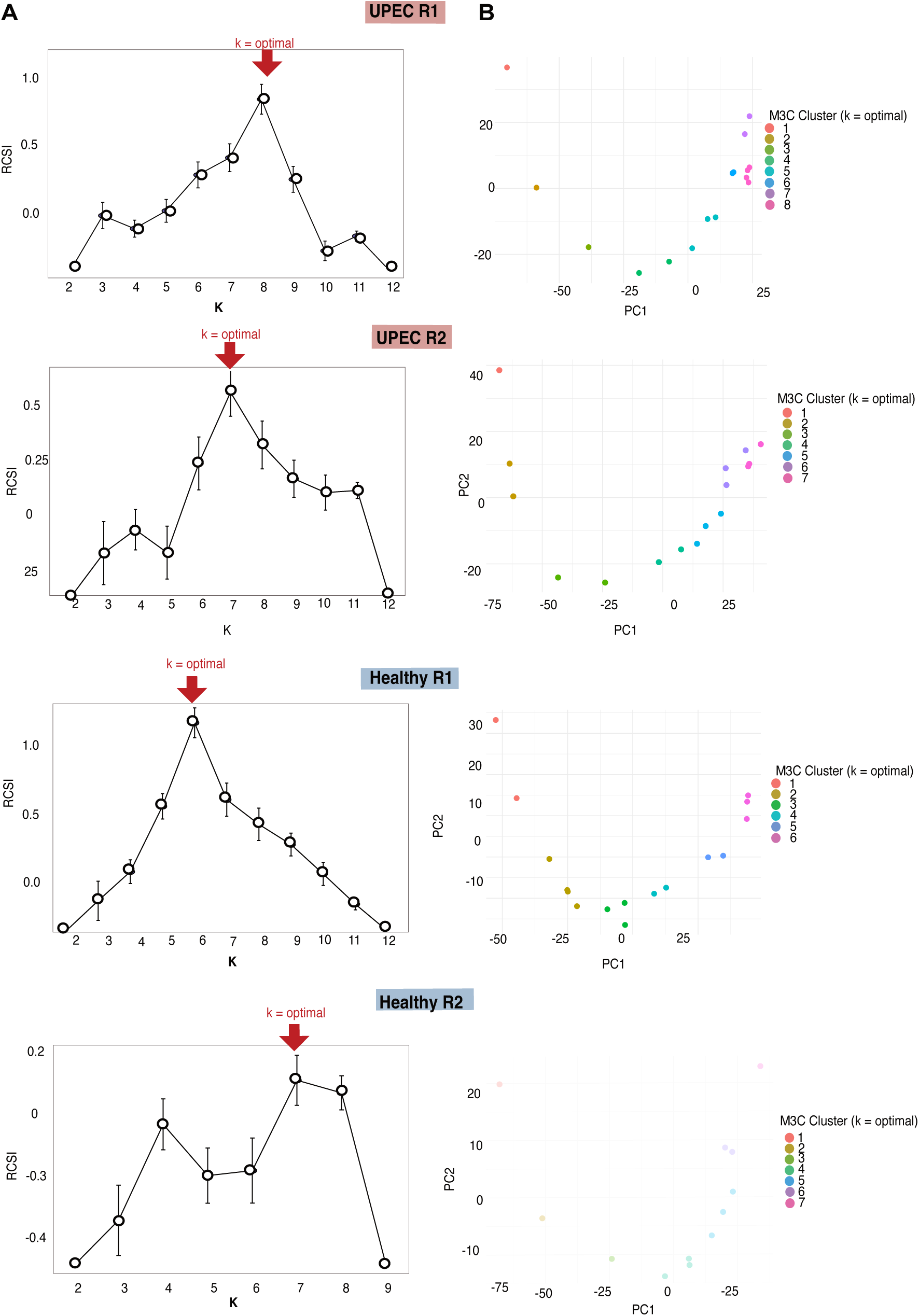
Cluster stability index of NIRL 3D proteomics data. **(A)** Relative cluster stability index (RCSI) from Monte Carlo Reference-based consensus clustering (M3C) for k=2 to k=12, individually calculated for each animal in the NIRL 3D proteomics data. The ideal cluster number is highlighted. **(B)** Scatter plot visualization of the first two principal components (PCs) from unsupervised PCA, based on all proteins (Healthy R1: N =4036; Healthy R2: N=4078; UPEC R1: N=4010; UPEC R1: N=4085). Colours represent different clusters at k=optimal. For PC calculation log2 transformed, normalized, and imputed values were used.

**Figure S3.**
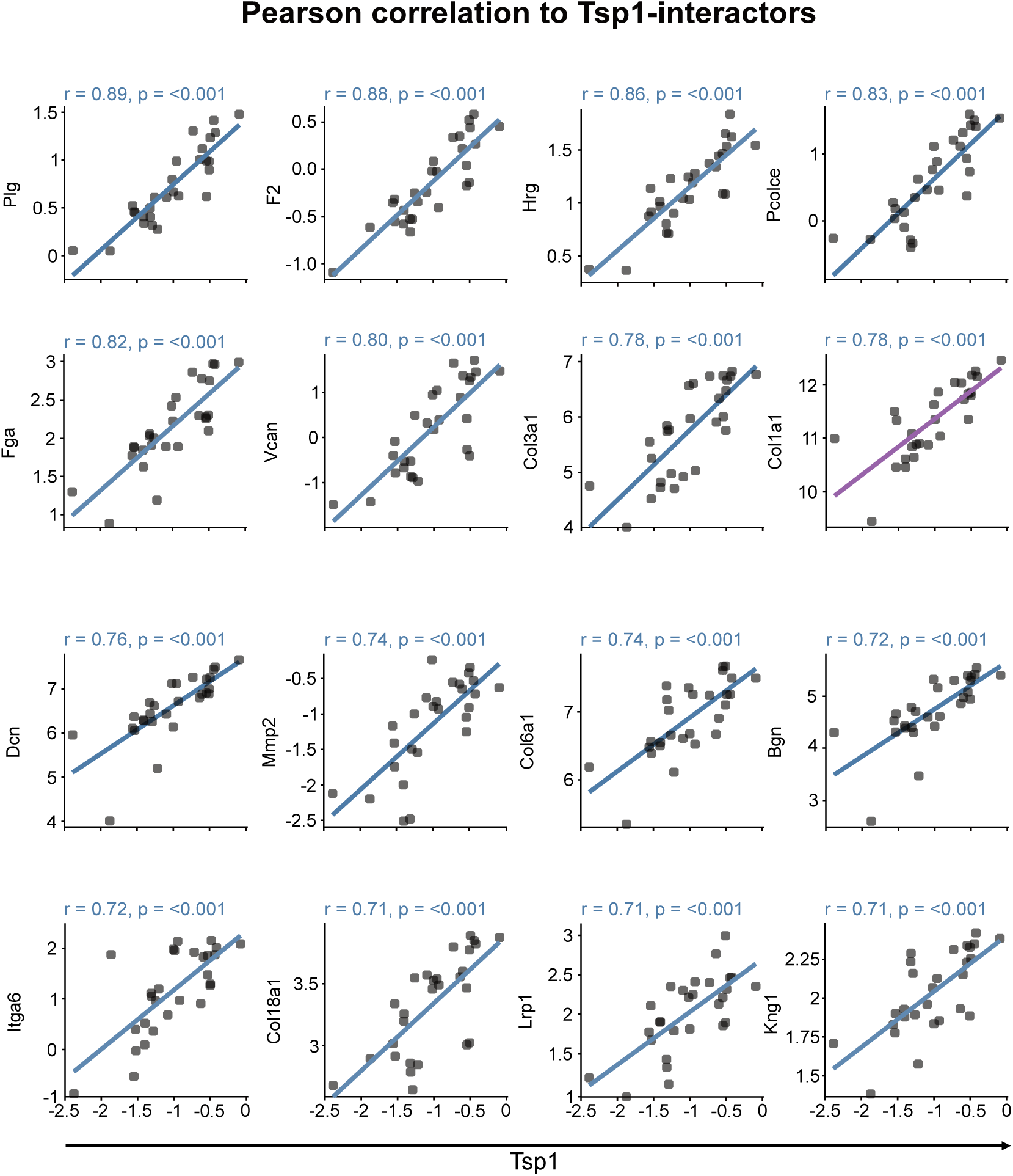
Correlation of Tsp1 with known interactors. Pearson correlation between Tsp1and known Tsp1 interactors^53^ in healthy animals. Only proteins with a strong linear correlation (> 70%) were considered. Proteins additionally identified as correlating with Tsp1 in UPEC bladders (r > 0.7) are highlighted. The Pearson correlation coefficient was calculated on log2 transformed, normalized, and imputed values. Scaling was not applied prior to correlation.

**Figure S4.**
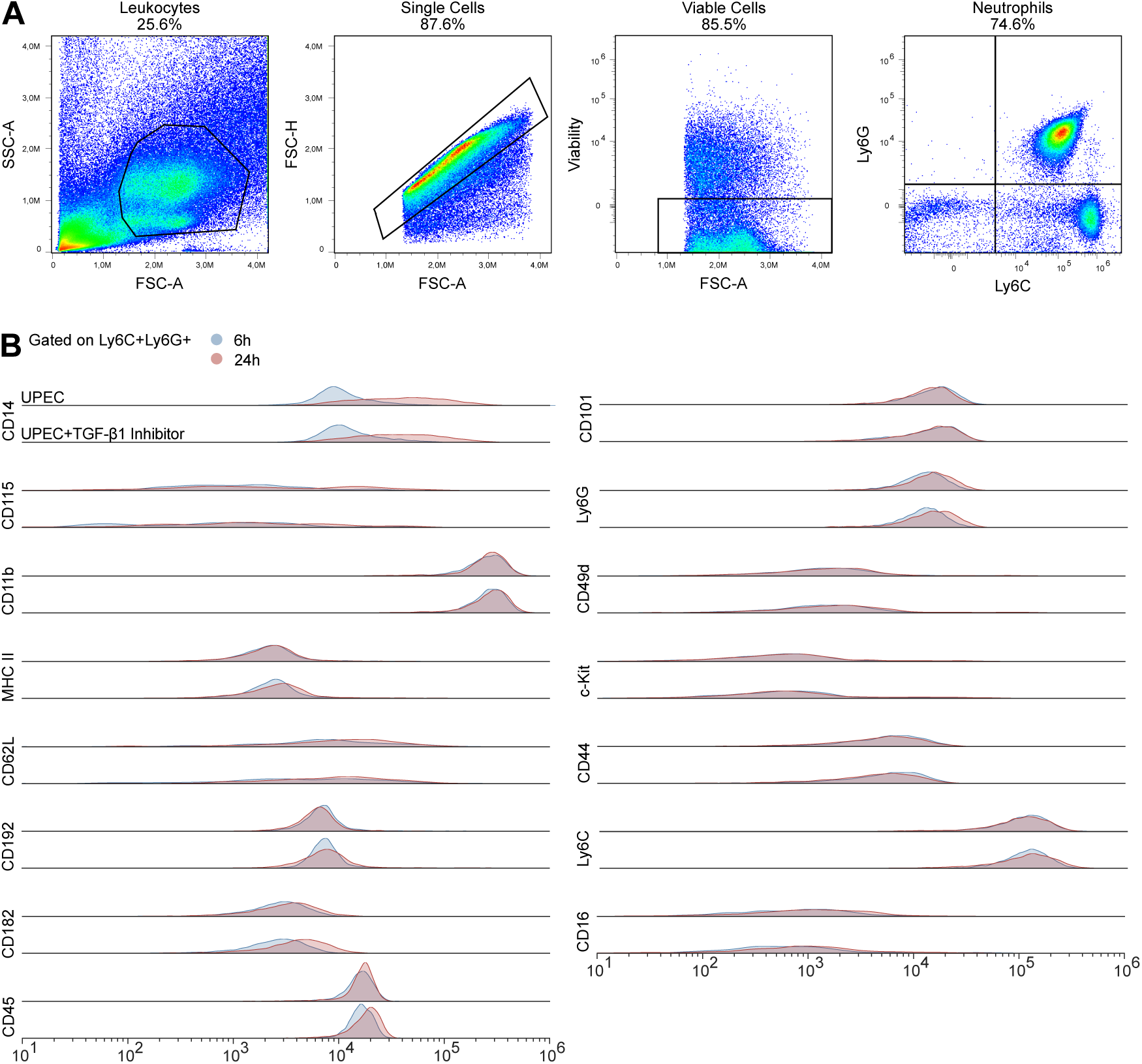
Maturation of neutrophils does not depend on TGF-β1. **(A)** Gating strategy for flow cytometric identification of neutrophils (Ly6G⁺Ly6C^+^) from bladder tissue 6 and 24 hours after infection. **(B)** Expression of surface markers on bladder neutrophils analysed at 6 hour (blue) and 24 hour (red) post-infection. Histograms show the density distribution of fluorescence intensity, allowing comparison of marker expression between time points and experimental groups. n=6 (6 and 24 hours UPEC); n= 5/6 (6 and 24 hours UPEC +TGF-β1 inhibitor).

## Notes

### Competing Interest Statement

The authors have declared no competing interest.

